# Protein arginylation is regulated during SARS-CoV-2 infection

**DOI:** 10.1101/2021.11.02.466971

**Authors:** Janaina Macedo-da-Silva, Livia Rosa-Fernandes, Vinicius de Moraes Gomes, Verônica Feijoli Santiago, Catarina Maria Stanischesk Molnár, Bruno R. Barboza, Edmarcia Elisa de Sousa, Edison Luiz Durigon, Claudio R. F. Marinho, Carsten Wrenger, Suely Kazue Nagahashi Marie, Giuseppe Palmisano

## Abstract

In 2019, the world witnessed the onset of an unprecedented pandemic. In September 2021, the infection by SARS-CoV-2 had already been responsible for the death of more than 4 million people worldwide. Recently, we and other groups discovered that SARS-CoV-2 infection induces ER-stress and activation of unfolded protein response (UPR) pathway. The degradation of misfolded/unfolded proteins is an essential element of proteostasis and occurs mainly in lysosomes or proteasomes. The N-terminal arginylation of proteins is characterized as an inducer of ubiquitination and proteasomal degradation by the N-end rule pathway. Here we present, for the first time, data on the role of arginylation during SARS-CoV-2 infection. We studied the modulation of protein arginylation in Vero CCL-81 and Calu-3 cells infected after 2h, 6h, 12h, 24h, and 48h. A reanalysis of *in vivo* and *in vitro* public omics data combined with immunoblotting was performed to measure the levels of ATE1 and arginylated proteins. This regulation is seen specifically during infections by coronaviruses. We demonstrate that during SARS-CoV-2 infection there is an increase in the expression of the ATE1 enzyme associated with regulated levels of specific arginylated proteins. On the other hand, infected macrophages showed no ATE1 regulation. An important finding revealed that modulation of the N-end rule pathway differs between different types of infected cells. We also confirmed the potential of tannic acid to reduce viral load, and furthermore, to modulate ATE1 levels during infection. In addition, the arginylation inhibitor merbromin (MER) is also capable of both reducing viral load and reducing ATE1 levels. Taken together, these data show the importance of arginylation during the progression of SARS-CoV-2 infection and open the door for future studies that may unravel the role of ATE1 and its inhibitors in pathogen infection.

## Introduction

In 2019, the world witnessed the onset of an unprecedented pandemic^1^. Patients in the capital and largest city in China’s Hubei province, Wuhan, developed pneumonia associated with infection with a new type of coronavirus, called severe acute respiratory syndrome coronavirus 2 (SARS-CoV-2)^2,3^. Clinical symptoms presented by infected patients ranged from mild to severe and included nonspecific manifestations such as fever, cough, sore throat, respiratory failure, muscle damage, and death^2,4,5,6^. Although there had been a great worldwide mobilization to contain the exponential spread of SARS-CoV-2, and even considered less pathogenic than other coronaviruses^7,8^, in September 2021 the death of more than 4 million people in the world were due to the new coronavirus infection^9^ (https://covid19.who.int/). The search for effective treatments and measures to fight the pandemic has driven studies that try to elucidate the infectious mechanisms of SARS-CoV-2^10^. As it belongs to the *Coronaviridae* family, the new coronavirus has similarities in structure and pathogenicity with SARS-CoV, both being single-stranded positive RNA viruses (+ssRNA)^11^. However, differences in the structural Spike (S) glycoprotein were identified in the SARS-CoV-2 which attributed greater efficiency in its dissemination compared to other coronaviruses^12,13^.

During the replication cycle, SARS-CoV proteins use the host’s endoplasmic reticulum (ER) to induce the formation of double-membrane vesicles for RNA synthesis, followed by the assembly of virions in the ER-Golgi intermediate compartment by structural proteins^14, 15^. The intense use of the host’s ER during viral replication increases the stress in this compartment, resulting in the accumulation of misfolded proteins and activation of the unfolded protein response (UPR)^15,16^. Recently, we^17^ and other groups discovered that SARS-CoV-2 infection also induces ER-stress and may be associated with patient survival^17,18,19^. The UPR pathway is highly conserved and regulates important cellular events such as growth, defense, homeostasis, and cell survival^13, 20^. In addition, this pathway activation may also boost tissue repair processes. The degradation of misfolded/unfolded proteins is an essential element of proteostasis^21^ and occurs mainly in lysosomes or proteasomes, which degrade long-lived and short-lived proteins, respectively^22, 23, 24^.

Ubiquitination is a universal tagging for protein degradation and is recognized by the proteasome^25, 26^. The N-terminal arginylation of proteins is characterized as an inducer of ubiquitination and proteasomal degradation by the N-end rule pathway, as elucidated by Varshavsky et. al^27, 28^. A direct relationship between the half-life of a protein and its N-terminal residue has been demonstrated^27, 28^. While methionine promotes protein stability, other amino acids, including arginine, result in rapid degradation^27, 28^. Studies have demonstrated the arginylation of different proteins in oxidized cysteine residues or aspartic acid and glutamic acid exposed at the N-terminus^29, 30, 31, 32, 33^. N-terminal asparagine and glutamine are tertiary destabilizing residues, as they can undergo an enzymatic deamidation reaction and be converted into glutamic acid and aspartic acid, recognized as secondary sites^33^. Thus, arginylated proteins or protein fragments with tertiary and secondary residues exposed at the N-terminus are universally recognized as marked for degradation through the ubiquitination-proteasome pathway^34, 35^. Moreover, internal aspartic and glutamic acids have been found to be sites specifically arginylated on several proteins involved in different biological processes^32, 36, 37^.

Protein arginylation has been associated with cellular stress conditions^38^, including ER-stress, oxidative stress, and misfolded protein stress^38, 39, 40^. Under these conditions, arginylation promotes cell death or growth arrest^40^. Furthermore, the activity of arginyl-tRNA-protein transferase (ATE1), the enzyme that promotes arginylation, is necessary to decrease mutation events when a cell is subjected to stressful conditions that damage DNA^39^. In 2002, the knockout of the ATE1 gene resulted in abnormalities in essential processes such as cardiac development, angiogenesis, and tissue morphogenesis in mammals^41^. However, the role of arginylation in infectious diseases has been little explored and is currently unknown.

In order to increase the knowledge on this topic, we present for the first-time data on the role of arginylation during SARS-CoV-2 infection. We conducted a study on the modulation of the N-end rule pathway and protein arginylation in Vero CCL-81 and Calu-3 cells infected after 2h, 6h, 12h, 24h, and 48h. A reanalysis of public omics data combined with western blotting was performed to measure the levels of ATE1 and arginylated proteins. We demonstrated that during viral infection there was an increase in the expression of the ATE1. Furthermore, there was a strong correlation between ATE1 levels and ER-related processes. We also demonstrated the potential of tannic acid and merbromin (MER) to reduce viral load, and furthermore, to modulate ATE1 levels during infection.

## Methods

### 1. Data sources and curation

Previously published studies were used to verify the abundance of proteins that make up the N-end rule pathway in non-infected and SARS-CoV-2 infected groups: (i) Saccon et al^42^ (Calu-3, Caco-2, Huh7, and 293FT cell lines, proteomics); (ii) Nie et al^43^ (autopsy 7 organs, 19 patients, proteomics); (iii) Leng et al^44^ (lung tissue, 2 patients, proteomics); (iv) Qiu et al^45^ (lung tissue, 3 patients, proteomics); (v) Bojkova et al^46^ (Caco-2 cells, proteomics); (vi) Wu et al^47^ (lung tissue, colonic transcriptomics); and (vii) Desai et al^48^ (lung tissue, transcriptomics). To verify modulation of the N-end rule pathway in other viral infections, including MERS-CoV/SARS-CoV/H1N1 influenza virus/Respiratory syncytial virus (RSV), data from the following studies were evaluated: (viii) Zhuravlev et al^49^ (MRC-5, A549, HEK293FT, and WI-38 VA-13 cell lines, H1N1 influenza virus, transcriptomics), (ix) Li et al^50^ (A549 and 293T cell lines, H1N1 influenza virus, transcriptomics), (x) Krishnamoorthy et al^51^ (comparative among coronaviruses, transcriptomics), (xi) Ampuero et al^52^ (time course of RSV infection in the lung, transcriptomics), (xii) Besteman et al^53^ (RSV infected neutrophils, transcriptomics), and (xiii) Dave et al^54^ (RSV infected alveolar cell, proteomics). Deep proteome data from non-infected cell lineages were recently publicly available by Zecha et al^55^ to model SARS-CoV-2 infection in Vero E6 (Kidney epithelial cell, African green monkey), Calu-3 (lung adenocarcinoma), Caco-2 (colorectal adenocarcinoma), and ACE2-A549 (lung carcinoma expressing ACE2 to gain cellular entry). The iBAQ intensities of proteins that make up the N-end rule pathway were evaluated without infection to accesses the basal levels of arginylation-related proteins. Experimentally arginylated proteins were retrieved from Seo et al^56^ and Wong et al^57^ datasets to access the regulation levels of these proteins during SARS-CoV-2 infection. Single-cell RNA-seq data from nasopharyngeal samples provided by Chua et al^58^ were reanalyzed to identify cell clusters expressing the ATE1 enzyme.

### 2. Bioinformatics analysis

The tidyverse^59^, biostrings, and seqinr^60^ packages were used to map potentially arginylated proteins in the *Homo sapiens* and *Chlorocebus sabaeus* proteomes (downloaded in May 2021, https://www.uniprot.org/). Signal peptide sequences were removed. Only proteins that have the potential to be arginylated at the N-terminus (NtE, NtD, NtC, NtN, NtQ) were retained. The corrplot package was used to evaluate the correlation between proteins/genes, applying a Spearman test with a cut-off significance of p-value < 0.05. Proteins subcellular locations were determined by UniProt release 12.4 (https://www.uniprot.org/news/2007/10/23/release) and pRoloc package^61^. The analysis of gene ontology (GO) was determined by the g:profile^62^ and DAVID^63^ tools. A q-value threshold of 0.05 was used, corrected by the Benjamini-Hochberg method^64^. InteractiVenn was used to build the Venn diagrams^65^. The String database v.11.5 was applied for protein network analysis (https://string-db.org/) with the following parameters: medium confidence score (0.400); textmining, co-expression, and neighborhood enabled.

### 3. Single-cell RNA-seq re-analysis

Expression matrices were loaded into RStudio (v. 4.0.3) with the Seurat package^66^. A filter to remove cells with less than 200 expressed genes or more than 25% of mitochondrial transcripts was applied using the ‘subset()’ function in each sample. Then cell counts were log-normalized by a size factor of 10,000 RNA counts and feature selection was performed by selecting the 2,000 genes with the highest dispersion. Unsupervised identification of anchor correspondences between the canonical correlation analysis (CCA) space of each sample normalized data was performed with the ‘FindIntegrationAnchors()’ function with 30 dimensions. After that, the data was integrated by ‘IntegrateData()’ function and scaled using ‘ScaleData()’. Principal component analyzes (PCA) and uniform approximation and projection dimension reduction (UMAP) with 30 principal components were applied. A nearest neighbor plot using 30 PCA reduction dimensions was calculated using ‘FindNeighbors()’, followed by clustering using ‘FindClusters()’ with a resolution of 0.5. The Metaboanalyst platform^67^ was used to evaluate differently regulated genes between cell clusters identified in the single-cell RNA-seq analysis.

### 4. Cell culture

Vero CCL-81 cells were cultured in DMEM medium supplemented with 10% fetal bovine serum (FBS), 100 U/ml penicillin-streptomycin, 4.5 g/L glucose, 2 mM L-glutamine, 1 mM sodium pyruvate, and 1.5 g/L NaHCO_3_. Calu-3 cells were cultured in DMEM medium supplemented with 20% FBS, 1% non-essential amino acids, 4.5 g/L glucose, 2 mM L - glutamine, 1 mM sodium pyruvate, 100 U/ml penicillin-streptomycin and 1.5 g/L NaHCO_3_. THP-1 cells were cultured in RPMI-1640 supplemented with 10% FBS, and 1% penicillin-streptomycin at 37 °C. All cells were kept in a humidified 5% CO_2_ atmosphere. The THP-1 monocytes cells were differentiated into macrophage-like cells as described by Gatto et al^68^ with few modifications. The THP-1 monocytes were induced to differentiate into macrophages by the addition of phorbol-12-myristate 13-acetate (PMA, 50 ng/mL) (ab120297, Abcam, UK) for 48 hours (h). After this time, the PMA-containing medium was replaced with fresh medium without PMA for 24 hours prior to SARS-CoV-2 infection. Cell differentiation was verified by microscopy evaluating cell spreading and adhesion.

### 5. Viral infection and quantification

In this study, SARS-CoV-2 isolate HIAE-02: SARS-CoV-2/SP02/human/2020/BRA (GenBank accession number MT126808)^69^ was used in all infections with multiplicity of infection (MOI) of 0.02. For comprehensive time course evaluation, Vero CCL-81 and Calu-3 cells were infected with SARS-CoV-2. Following adsorption in DMEM with 2.5% FBS for 1h, fresh medium was added, and cells were further incubated at 37 °C and 5% CO_2_. Cell lysates were collected at 2, 6, 12, 24, and 48h post infection (hpi) in 8M urea supplemented with protease (cOmplete, Sigma-Aldrich) and phosphatase inhibitors (PhosStop, Sigma-Aldrich). Aliquots of cells and supernatants were collected at the different time points for virus RNA copy number quantification by reverse transcription-quantitative polymerase chain reaction (RT-qPCR), targeting the E gene as previously described^70^.

To evaluate the effects of protein arginylation inhibition, Calu-3 cells and differentiated macrophages were incubated with 25μM of merbromin (Mercury dibromofluorescein disodium salt, Sigma-Aldrich), 1μM of tannic Acid (Sigma-Aldrich) or medium for 1h prior to infection with SARS-CoV-2. Following adsorption in DMEM with 2.5% FBS for 1h, infected and respective mock infected cells were kept at the same inhibitor concentration for 24 and 72 hpi at 37 °C and 5% CO_2_. Cell lysates were collected in BE buffer (HEPES 10mM, SDS 1%, MgCl2.6H2O 1,5mM, KCl 10mM, DTT 1mM, NP-40 0,1%) containing protease (cOmplete, Sigma-Aldrich) and phosphatase inhibitors (PhosStop, Sigma-Aldrich). Aliquots of supernatants were collected at the different conditions for RNA extraction using TRIzol reagent (ThermoFisher) according to the manufacturer’s instructions. Viral copy number quantification by RT-qPCR was performed using Detection Kit for 2019 Novel Coronavirus (2019-nCoV) RNA (PCR-Fluorescent Probing) (Cat. #DA-930) (China) in a QuantStudio 3 real-time PCR system (Applied Biosystems) according to the manufacturer’s instructions. The percentage of viral release was calculated with the CTs values of the experimental triplicates. Graphics were done using GraphPad Prism software version 8.1 (GraphPad Software, San Diego, USA).

All assays were conducted in biological triplicates in a BSL-3 facility at the Institute of Biomedical Sciences, University of São Paulo, under the Laboratory biosafety guidance related to coronavirus disease (COVID-19): Interim guidance, 28 January 2021 (https://www.who.int/publications/i/item/WHO-WPE-GIH-2021.1).

### 6. Western blotting

Proteins were extracted from cellular lysates and quantified using the Qubit Protein Assay Kit platform (Invitrogen) according to the manufacturer’s instructions. A total of 15μg of proteins were separated by SDS-PAGE and electrotransferred to PVDF membranes, which were directly incubated with blocking buffer (5% bovine serum albumin (BSA) in Tris-buffered saline (TBS) at 0.05% Tween-20 (TBST) for 1h. Subsequently, samples were incubated with primary antibodies (**Table 1**) overnight and washed three times with TBST. Then, the bands were incubated with the respective secondary antibodies for 1h at room temperature. Immunoreactive bands were detected with the ChemiDoc XRS Imaging System equipment and protein quantification was performed using the ImageJ software. Graphs were plotted using GraphPad Prism version 8.1 software. Bands with statistically significant intensities among groups were evaluated by applying an Ordinary One-way ANOVA, with Tukey post-hoc test (0.05 cut-off).

**Table 1.**
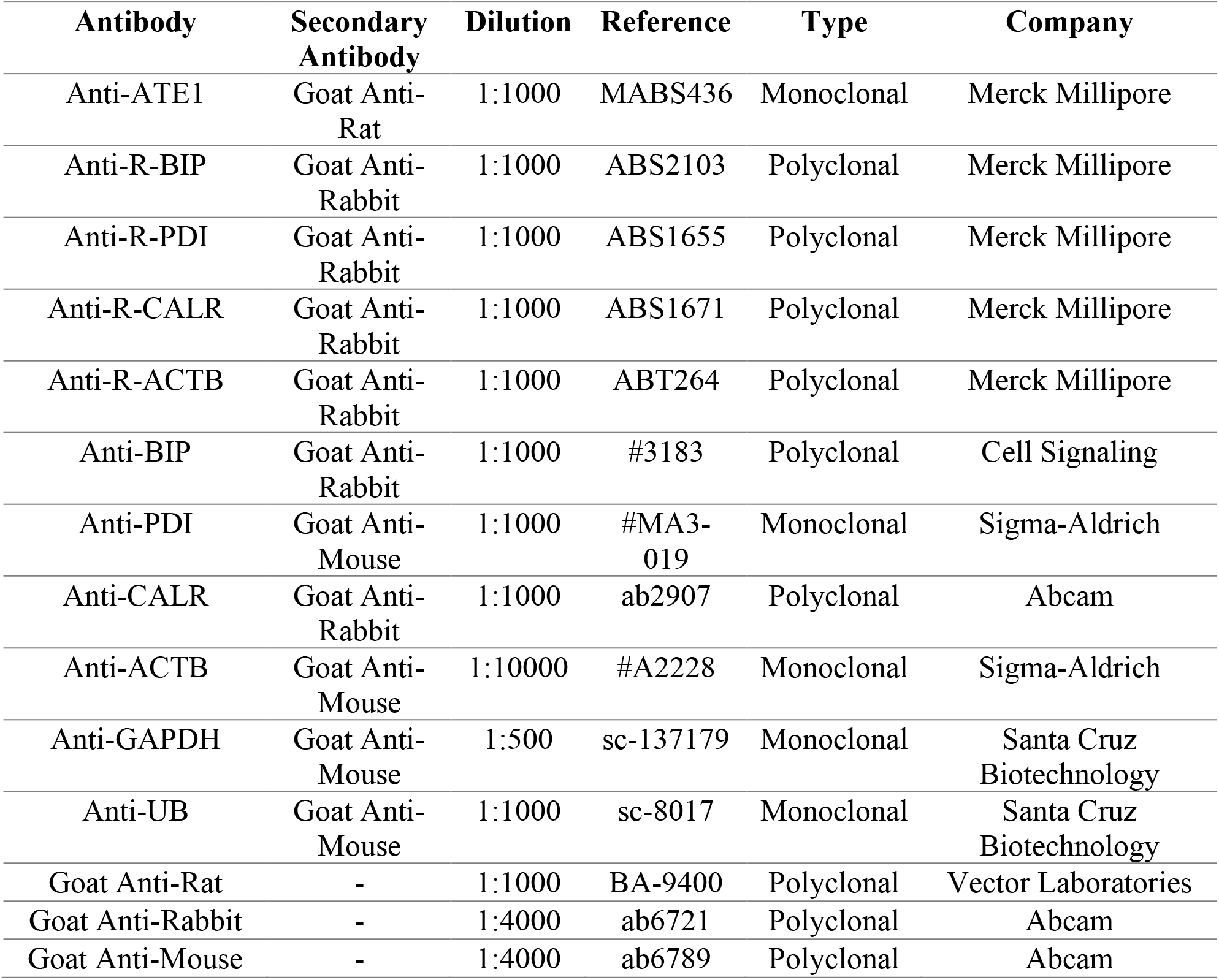
Primary and secondary antibodies used in western blotting analyses, with their respective dilutions, reference catalog number, type, and Supplier Company.

| **Antibody** | **Secondary Antibody** | **Dilution** | **Reference** | **Type** | **Company** |
| --- | --- | --- | --- | --- | --- |
| Anti-ATE1 | Goat Anti-Rat | 1:1000 | MABS436 | Monoclonal | Merck Millipore |
| Anti-R-BIP | Goat Anti-Rabbit | 1:1000 | ABS2103 | Polyclonal | Merck Millipore |
| Anti-R-PDI | Goat Anti-Rabbit | 1:1000 | ABS1655 | Polyclonal | Merck Millipore |
| Anti-R-CALR | Goat Anti-Rabbit | 1:1000 | ABS1671 | Polyclonal | Merck Millipore |
| Anti-R-ACTB | Goat Anti-Rabbit | 1:1000 | ABT264 | Polyclonal | Merck Millipore |
| Anti-BIP | Goat Anti-Rabbit | 1:1000 | #3183 | Polyclonal | Cell Signaling |
| Anti-PDI | Goat Anti-Mouse | 1:1000 | #MA3-019 | Monoclonal | Sigma-Aldrich |
| Anti-CALR | Goat Anti-Rabbit | 1:1000 | ab2907 | Polyclonal | Abcam |
| Anti-ACTB | Goat Anti-Mouse | 1:10000 | #A2228 | Monoclonal | Sigma-Aldrich |
| Anti-GAPDH | Goat Anti-Mouse | 1:500 | sc-137179 | Monoclonal | Santa Cruz Biotechnology |
| Anti-UB | Goat Anti-Mouse | 1:1000 | sc-8017 | Monoclonal | Santa Cruz Biotechnology |
| Goat Anti-Rat | - | 1:1000 | BA-9400 | Polyclonal | Vector Laboratories |
| Goat Anti-Rabbit | - | 1:4000 | ab6721 | Polyclonal | Abcam |
| Goat Anti-Mouse | - | 1:4000 | ab6789 | Polyclonal | Abcam |

## Results

### SARS-CoV-2 infection modulated the N-end rule pathway and increased ATE1 enzyme expression

To explore protein arginylation during SARS-CoV-2 infection, we performed a study combining multi-omics (*in silico*) analysis and validated the findings mined *in silico* in time-course SARS-CoV-2 infection experiments at protein level by immunoblotting (**Fig. 1A**). Initially, the basal levels of enzymes involved in the N-end rule pathway were evaluated in different uninfected cells and compared to the total proteome profile (**Fig. 1B**). Enzymes involved in protein arginylation (ATE1), ubiquitination (UBR1, UBR2, UBR4, UBR5), arginine-tRNA ligase assembly (RARS2), deamidation (NTAN1), and N-terminal methionine removal (CASP6, CASP7, CASP8, CASP9, METAP1, METAP2, CASP10, CASP2, CASP3, CAPN7, CAPN1, CAPN2, CAPN5) were identified in all cell models with no statistical difference among them. These findings indicated that enzymes involved in the protein arginylation pathway were not modulated based on cell type or on specie in uninfected conditions. A total of 1,152 proteins (**Fig. 1C**) have the potential to be arginylated at the N-terminus (NtE, NtD, NtC, NtN, NtQ), in agreement with the UniProt sequence. These proteins were identified in uninfected Calu-3, Vero E6, Caco-2, and ACE2-A549 cell lines showing a similar expression pattern (**Fig. 1D**), regardless of the organism (Green Monkey and Human).

**Figure 1.**
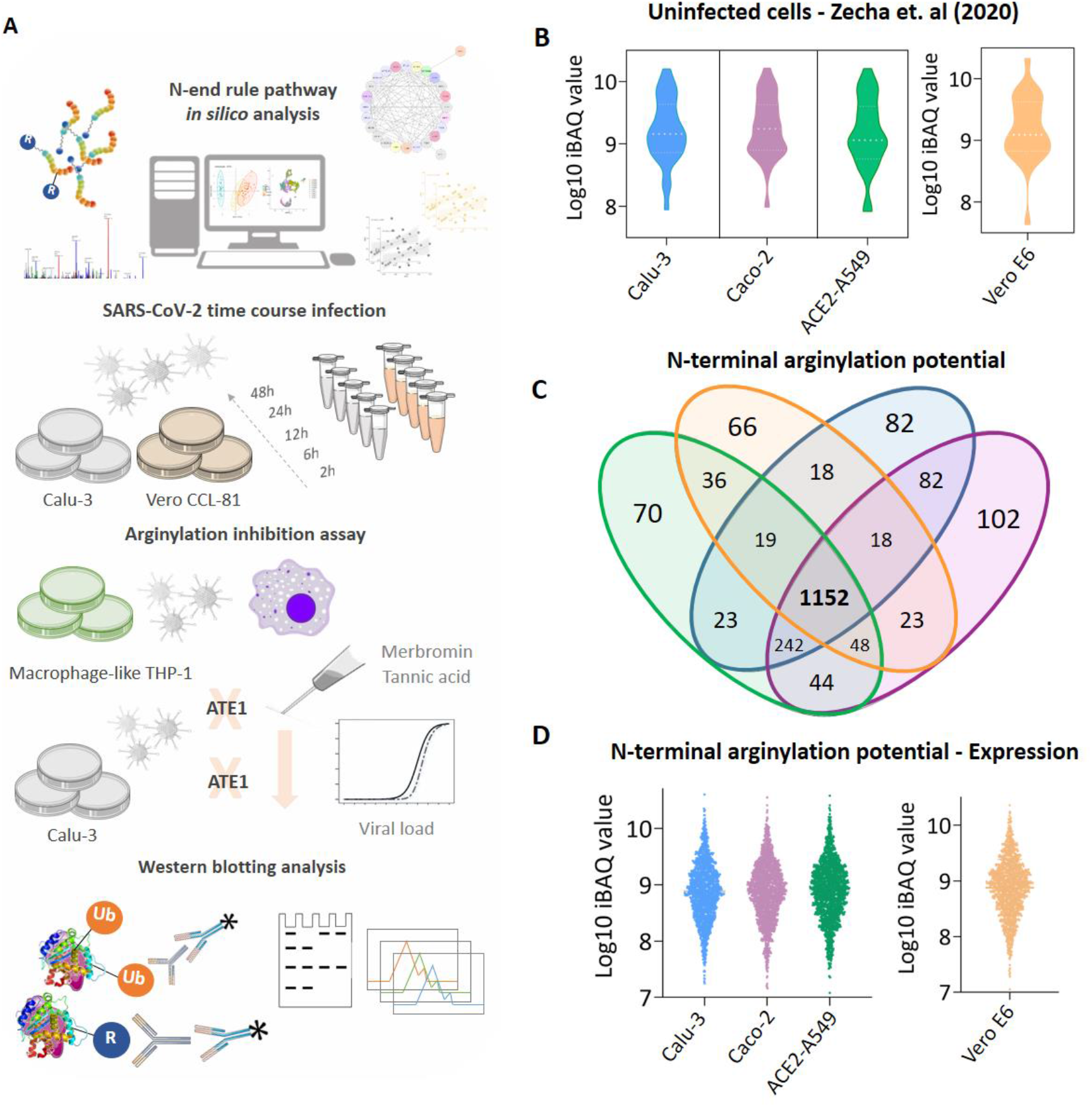
N-end rule pathway modulation in uninfected cell models. (**A**) Experimental workflow adopted to identify the modulation of protein arginylation during SARS-CoV-2 infection; (**B**) Expression profile of enzymes participating in the N-end rule pathway identified in uninfected Calu-3 (blue), Caco-2 (purple), ACE2-A549 (green), and Vero E6 (orange) cells; (**C**) Proteins with potential to be arginylated at N-terminal (NtE, NtD, NtC, NtN, NtQ) identified and quantified in uninfected cell models; (**D**) Expression profile of the 1,152 proteins with potential to be arginylated identified in uninfected Calu-3, Caco-2, ACE2-A549, and Vero E6 cells. Proteins with the potential to be arginylated were determined based on their sequences deposited in Uniprot (*Homo sapiens* and *Chlorocebus sabaeus*).

Furthermore, we reanalyzed 8 datasets covering transcriptomic and proteomic data of *in vitro* and *in vivo* SARS-CoV-2 infection of different biological systems^42, 43, 44, 45, 46, 47, 48^ (**Fig. 2A**). ATE1 expression was higher during infection most of the datasets, being significantly upregulated at both transcript and protein level ^42, 45, 46, 47, 48^. On the other hand, RARS1 and RARS2 protein expressions were opposite, being RARS1 upregulated and RARS2 (mitochondrial) downregulated. UBR1, UBR2, and UBR5 ubiquitin-ligases (E3) expressions were increased in infection; however, UBR4 was regulated in different directions at transcript (Wu et al) ^47^ and protein (Saccon et al) ^42^ levels. The expression of proteins involved in the removal of N-terminal methionine were variable among the different studies. However, the protein expressions of the caspase family were upregulated, especially the CASP3 expression was statistically significant in four studies^42, 44, 45, 47^. To confirm the above listed findings, western blotting analysis was performed to measure ATE1 levels (**Fig. 2B**). In agreement with the omics data, SARS-CoV-2 infected Calu-3 and Vero CCL-81 cells had statistically higher ATE1 levels compared to the CTRL uninfected group (**Fig. 2B**). Time-course data revealed an increase from 2h onwards in ATE1 in Vero CCL-81 cells; on the other hand, in Calu-3 cells statistical significance between the groups was found only after 48h (p-value = 0.0259), possibly due to the less susceptibility of these cells to SARS-CoV-2 infection. Both cell lines showed an equilibrium trend in ATE1 levels after 6h, however, higher levels were observed in Calu-3 cells. It was verified that E2 ubiquitin-conjugating enzymes (UBE2G2, UBE2L3, UBE2D2, UBE2D3, UBE2K, UBE2D4, UBE2R2, UBA52, UBE2A, UBA3, UBE2W, UBE2L6, and UBE2E1) were also upregulated in the infected groups (**Supplementary File 1**). As with the omic data, there was an increase in ubiquitinated proteins confirmed by western blotting analysis in Vero CCL-81 cells (**Fig. S1A**), revealing that 48h after the onset of infection there was an increase in this event. On the other hand, in Calu-3 cells, the assay did not identify ubiquitinated proteins, possibly due to less permissive environment towards infection and slower protein arginylation kinetics at the infection conditions considered in this work (**Fig. S1B**).

**Figure 2.**
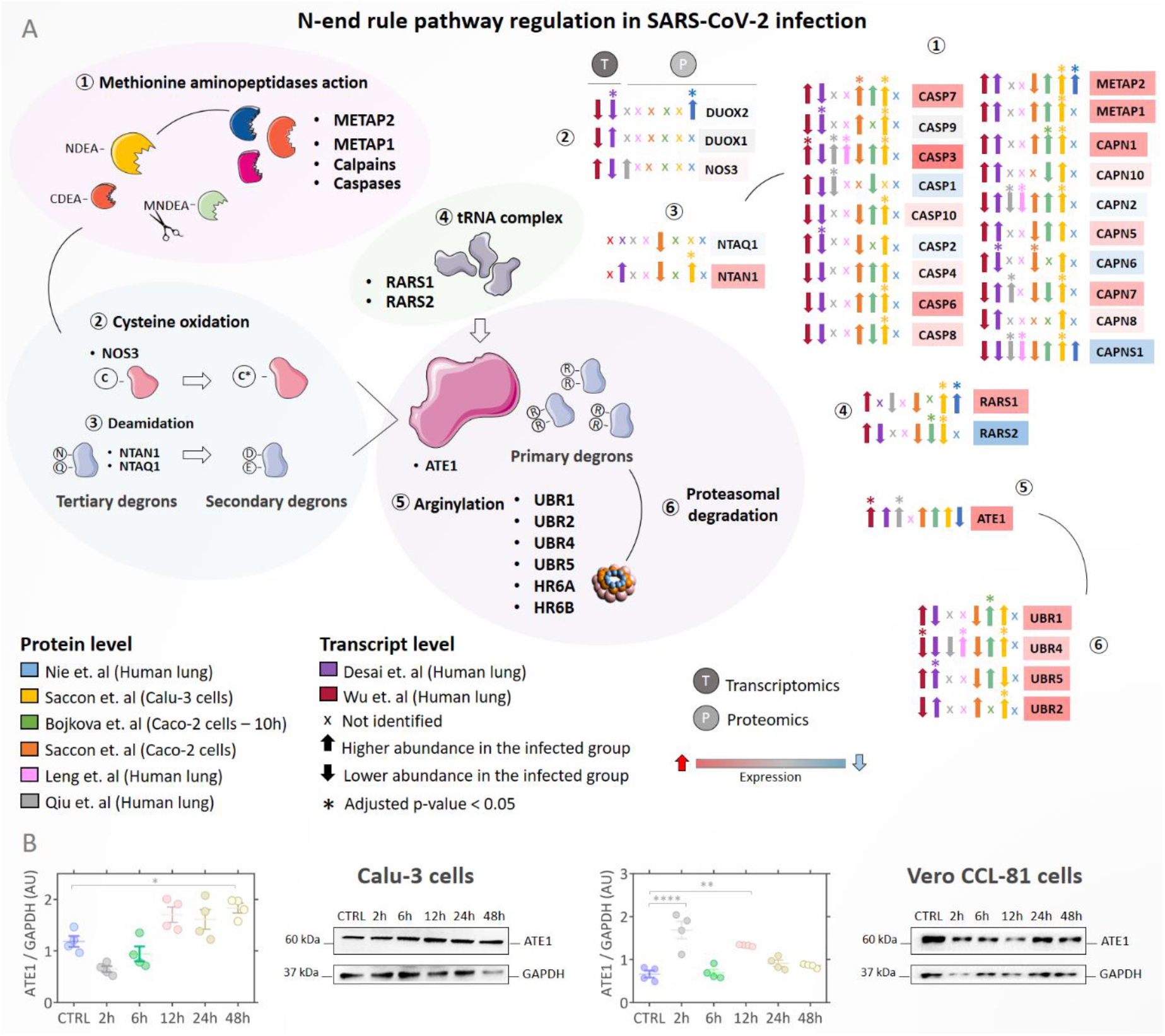
N-end rule pathway modulation in infected *in vitro* and *in vivo* models. (**A**) Regulation of proteins involved in the N-end rule pathway during SARS-Cov-2 infection. Proteins/genes were considered differentially regulated if they had a q-value < 0.05 (Benjamini-Hochberg) and are indicated by the symbol (*). Up arrows indicate proteins/genes with the higher abundance in the infected group (INF) and down arrows indicate proteins with lower abundance in the infected group. The color of the boxes proteins/genes indicates the fold change (INF/CTRL) considering all studies evaluated; the red color indicates higher abundance in the infected group and the blue color lower abundance in the INF group. The symbol (x) indicates that a protein/gene was not identified in the dataset; (**B**) Western blotting analysis of ATE1 protein in Calu-3 and Vero CCL-81 cells infected with SARS-CoV-2 (classical strain) after 2h, 6h, 12h, 24h, and 48h.

### Increased ATE1 expression in SARS-CoV-2 infection was correlated with events linked to the endoplasmic reticulum (ER)

Once the increased abundance of ATE1 was confirmed in the infection, a multi-correlation analysis was performed using omics data to verify which proteins correlated with ATE1 (**Fig. 3A**). Only differentially regulated proteins/genes were selected from six studies^42, 43, 45, 46, 47, 48^. A total of 365 proteins/genes presented a significant correlation (p-value < 0.05) with ATE1 in at least two studies and 28 in at least three studies (**Supplementary File 2**). Analyzing the molecular functions (MF) of the 28 correlated proteins/genes, the enrichment of processes related to unfolded protein binding, protein-folding chaperone, and ubiquitin-protein ligase binding were found (**Fig. 3B**). Among the biological processes (BP), events related to ER and viral infection were enriched, such as protein target to ER, protein localization to ER, viral gene expression, and viral transcription (**Fig. 3C**). Pathways related to alterations in processes linked to RNA and coronavirus infection were also enriched (**Fig. 3D**). The GBP2 protein, involved in cellular response to infections, was correlated with ATE1 in four studies. Due to the observed relationship between the processes linked to the ER (**Fig. 3B** and **C**), we monitored the direction of the correlation of HSPBP1 (**Fig. 3E**) and HSP90B1 (**Fig. 3F**) with ATE1. These proteins showed significant positive correlations, except for the negative correlation observed in lung tissue by Qiu et al^45^.

**Figure 3.**
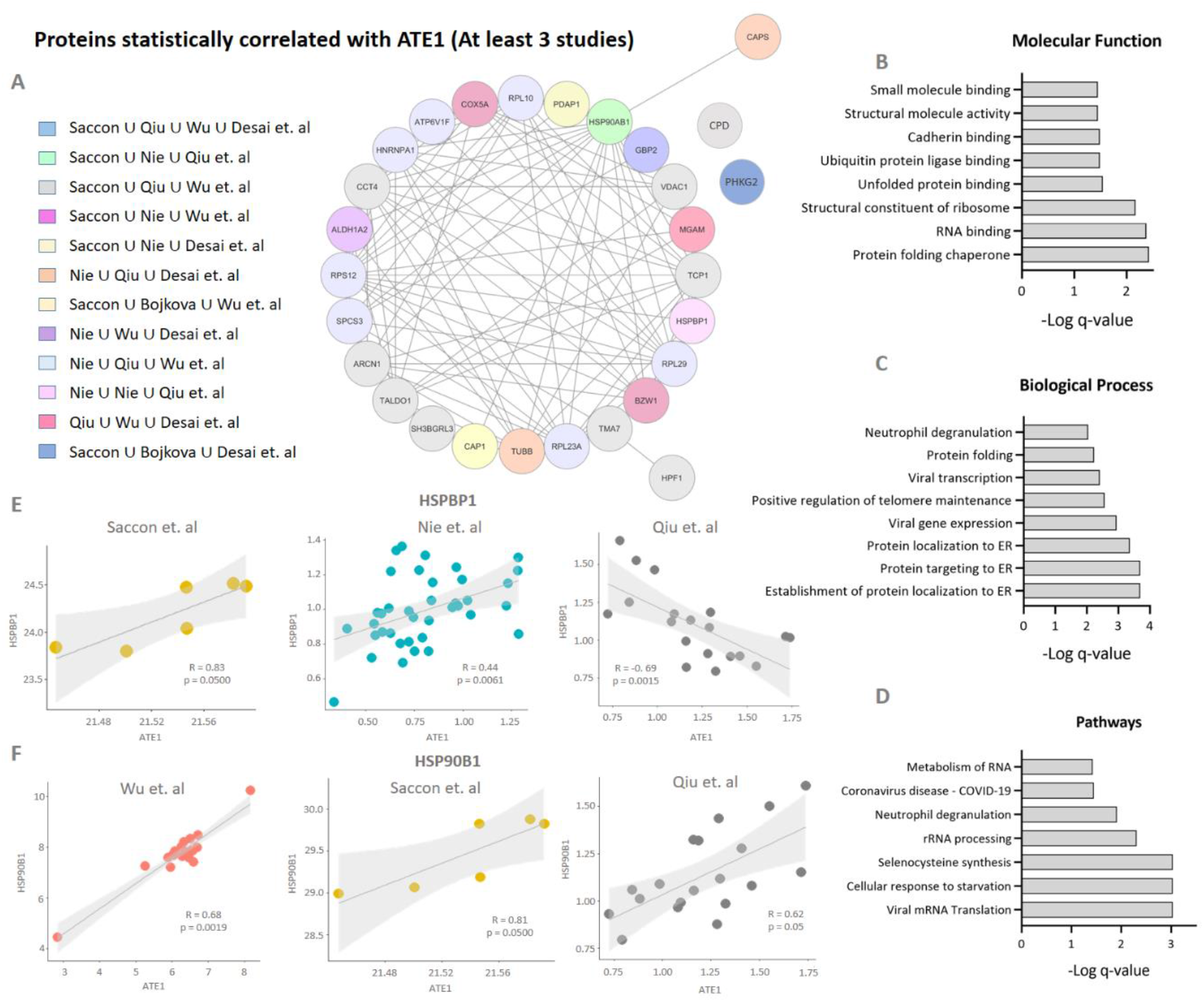
Multi-correlation expression analysis. **(A)** Proteins and genes correlated with ATE1 expression in at least three reanalyzed studies. The correlation analysis was determined by applying the Spearman test with a cut-off significance of p-value < 0.05. Only differentially regulated proteins/genes were considered to the correlation analysis; (**B**) Gene ontology (GO) analysis of molecular functions; (**C**) Biological processes and (**D**) pathways related to proteins/genes correlated with ATE1 in at least three studies; (**E**) Correlation graph of HSPBP1 and (**F**) HSP90AB1 proteins indicates the positive/negative correlation with ATE1.

After identifying a relationship between ATE1 and ER-associated chaperones/processes during SARS-CoV-2 infection, the arginylation levels of proteins located in the ER, heat shock protein family A (Hsp70) member 5 (HSPA5, also known as BIP), calreticulin (CALR), and protein disulfide isomerase (PDI) were analyzed by western blotting (**Fig. 4**). The BIP/HSPA5 arginylated protein level increased in both cell models over time with statistical significance 48h after the onset of infection (**Fig. 4A**). Interestingly, the arginylated CALR protein level decreased in Vero CCL-81 cells while increased in Calu-3 cells (**Fig. 4B**) compared to the CTRL uninfected cells. The PDI protein showed a significant decrease 6h after the start of infection in Calu-3 cells and is statistically more arginylated in Vero CCL-81 cells infected after 48h (**Fig. 4C**).

**Figure 4.**
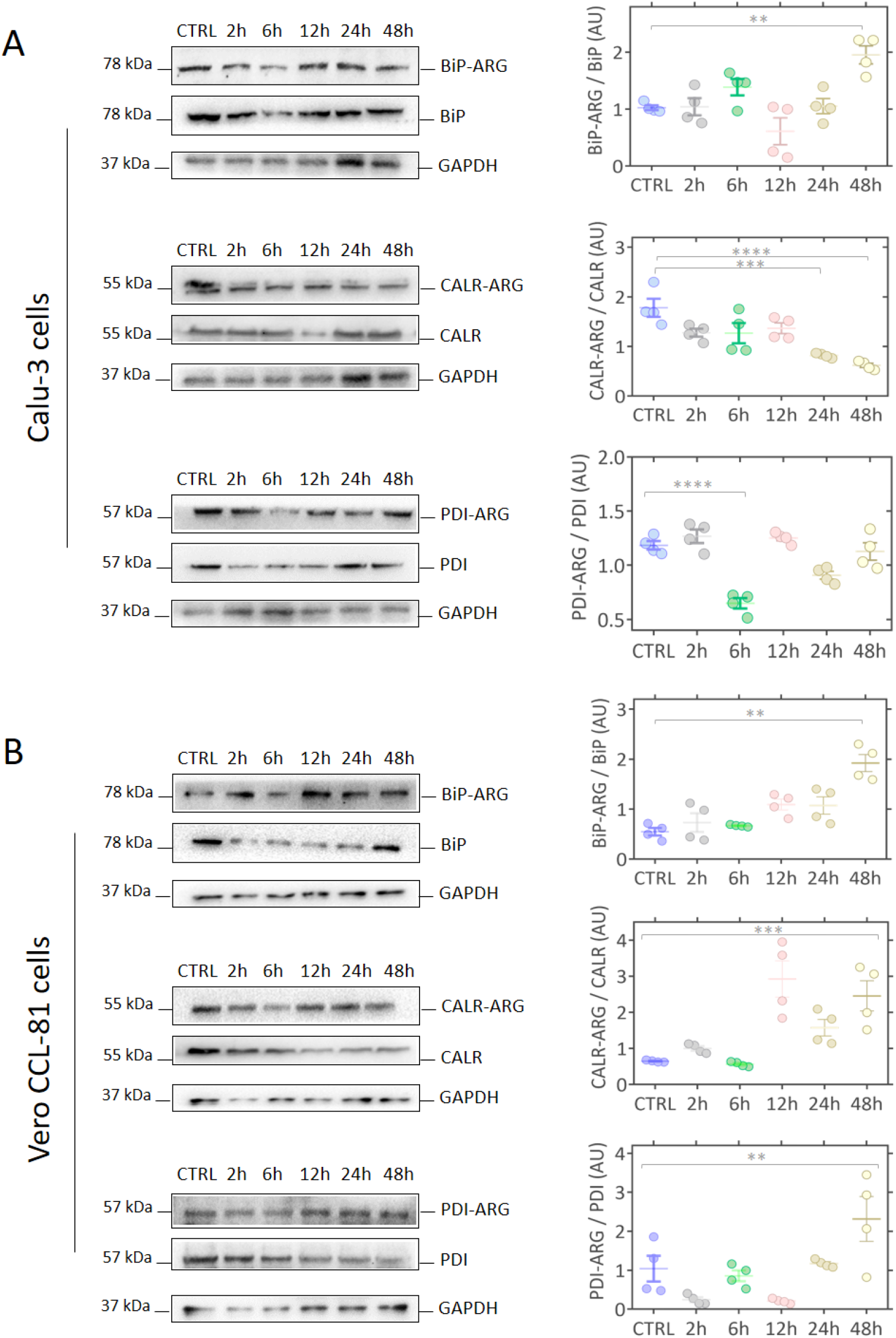
Modulation of arginylated proteins located in the endoplasmic reticulum (ER). (**A**) Representative western blot images of R-BiP/BiP, R-CALR/CALR and R-PDI/PDI proteins in Calu-3 and (**B**) Vero CCL-81 cells after 2h, 6h, 12h, 24h, and 48h of infection (classic strain). Each point represents an independent experiment (n = 3). The level of significance indicates: **** p <0.0001; *** p<0.001; ** p<0.005.

### Arginylation-related proteins were located mainly in the endoplasmic reticulum (ER) and cytoskeleton

Searching for other organelles involved in arginylation during SARS-CoV-2 infection, we performed a subcellular localization analysis of proteins correlated with ATE1 in at least two studies (**Fig. 5A**). These proteins mostly occupy complexes of chaperones, ribosomal, proteasome, and cytoskeletal microtubules and actin filament. Recently, Seo et al^56^ and Wong et al^57^ demonstrated experimentally that 152 were arginylated proteins, including mainly actins, chaperones, ribosomal components, and tubulins (**Supplementary File 3**), and nine proteins (VIM, HSPB1, PRDX4, ACTG1, ACTB, CALR, ATP5F1A, SPTAN1, and HSPA1B) overlapped in both studies. Bringing together arginylated proteins that were differentially regulated during SARS-CoV-2 infection and presented the same direction of regulation (upregulated or downregulated) in at least two studies (**Fig. 5B**), we observed that tubulins and chaperones were increased in the INF groups, on the other hand, VIM and SPTAN1 proteins were downregulated. Collectively, data analysis on differentially regulated proteins pointed to an increased level of arginylated proteins in SARS-CoV-2 infection (**Fig. 5B**). Looking at the arginylated proteins evaluated here (**Fig. 4**); we found that they are differentially regulated in different directions in the studies by Saccon et al^42^, Nie et al^43^, Wu et al^47^, and Leng et al^44^ (**Fig. 5B and C**). The subcellular location of the 152 arginylated proteins were mainly in the cytoskeleton, cytoplasm, and nucleus (**Fig. 5D**). Since the ACTB protein was previously identified as arginylated by Seo et al^56^ and Wong et al^57^, western blotting analysis was performed to measure arginylated ACTB levels in infected Vero CCL-81 and Calu-3 cells (**Fig. 5E and F)**. An increasing arginylation was observed up to 24h in Calu-3 cells, with a reduction 48h after infection. On the other hand, in Vero CCL-81 cells, the increase in arginylation occured only after 48h. Similarly, ACTB downregulation was identified by Wu et al in lung tissue obtained from patients who died of COVID-19 in Wuhan, China (**Supplementary File 1**).

**Figure 5.**
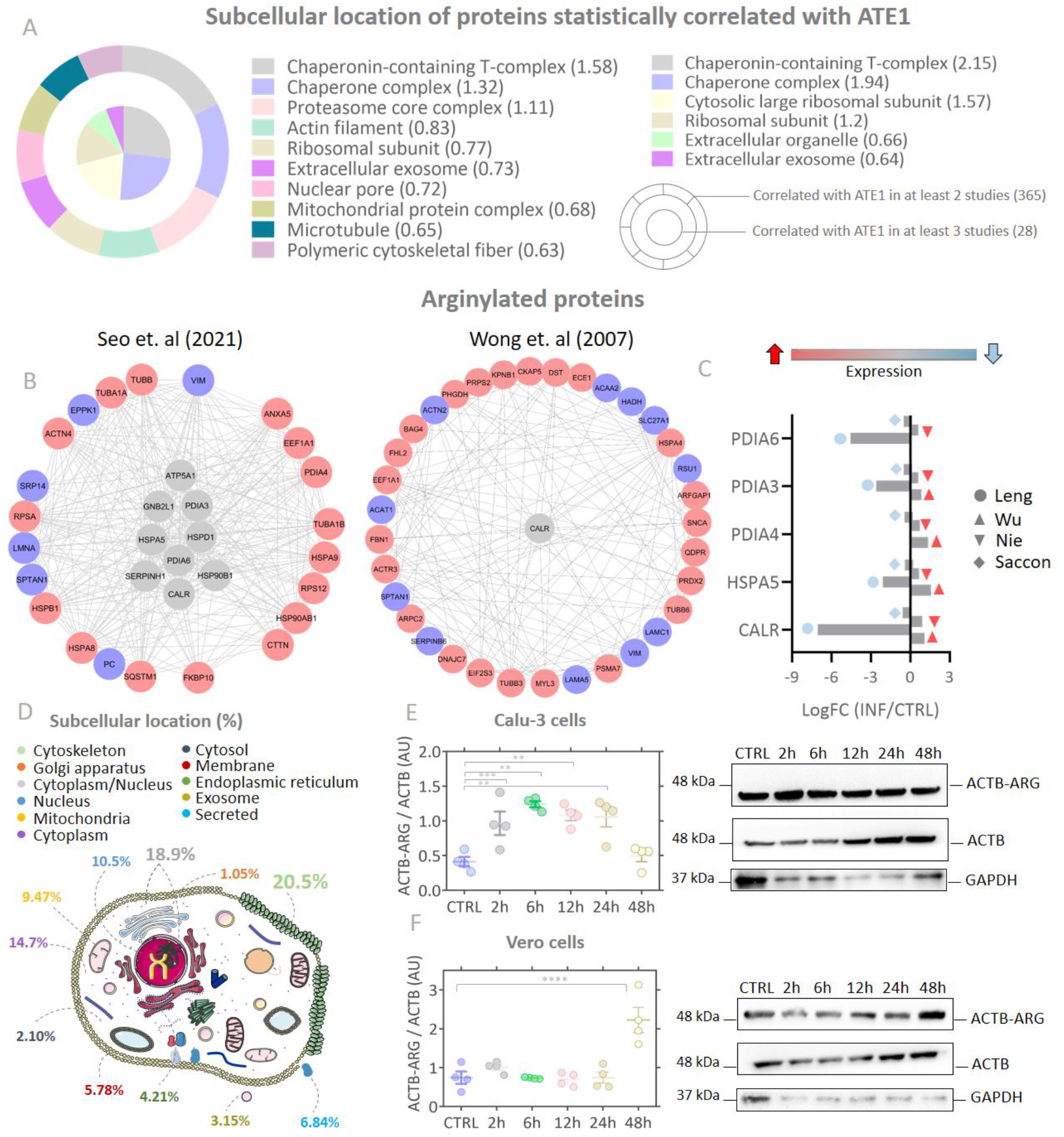
Subcellular localization of arginylation-related proteins. (**A**) Subcellular location of ATE1-correlated proteins in at least 2 studies (major circle) or in at least 3 studies (minor circle); (**B**) Arginylated proteins differently regulated during SARS-CoV-2 infection. The network shows proteins that were regulated at the same direction in at least two studies; (**C**) Arginylated endoplasmic reticulum (ER) chaperones regulation. The red and blue colors show up-regulated and down-regulated proteins, respectively, according to the foldchange (INF/CTRL). The gray color of the centered proteins indicates that the protein was differentially regulated in different directions in more than two studies; (D) Subcellular localization of arginylated proteins experimentally identified in Seo et al ^46^ and Wu et al ^57^; (E) Western blotting indicating arginylation of ACTB protein in Calu-3 cells and (F) Vero CCL-81 at 2, 6, 12, 24, 48h after infection.

### Tannic acid and merbromin reduced ATE1 expression level and viral load

In view of the close relationship between arginylation and SARS-CoV-2 infection demonstrated by the previous data, an enzyme inhibition assays by 1μM tannic acid and 25μM merbromin (MER) in Calu-3 cells were performed (**Fig. 6**). These concentrations of MER and tannic acid did not affect cellular viability. Notably, the infected cells before any treatment (INF-24h) presented higher expression of ATE1 than uninfected cells (CTRL), and the treatment with tannic acid and MER decreased significantly the ATE1 expression at 24 and 72h, respectively, to the level observed in uninfected (CTRL) cells. Interestingly, the expression level of the cytoskeleton protein, ACTB, also decreased after to the treatment with ATE1 inhibitors, corroborating the findings that the increased expression of ACTB after 24h after infection was associated to ATE1 activity (**Fig. 5E**). In addition, tannic acid inhibited arginylation in earlier time point (24h) compared to MER (72h). The ER proteins (CALR and PDI) expression levels decreased similarly to ATE1 and ACTB, however less pronounced, due to reduced arginylation after exposure to inhibitors. Interestingly, the BiP/HSP5A protein expression pattern differed from the other proteins (**Fig. 6**). There was an increase in the abundance of arginylated protein in relation to the CTRL group, which corroborated that the abundance of ATE1 was possibly not only related to the reduction in the levels of arginylation of the BiP/HSPA5 (**Fig. 4A**). Of note, tannic acid and MER were able to reduce the viral load or prevent virus entry into the cell. Such effect was more relevant in the inhibition with MER (**Fig. 6C**).

**Figure 6.**
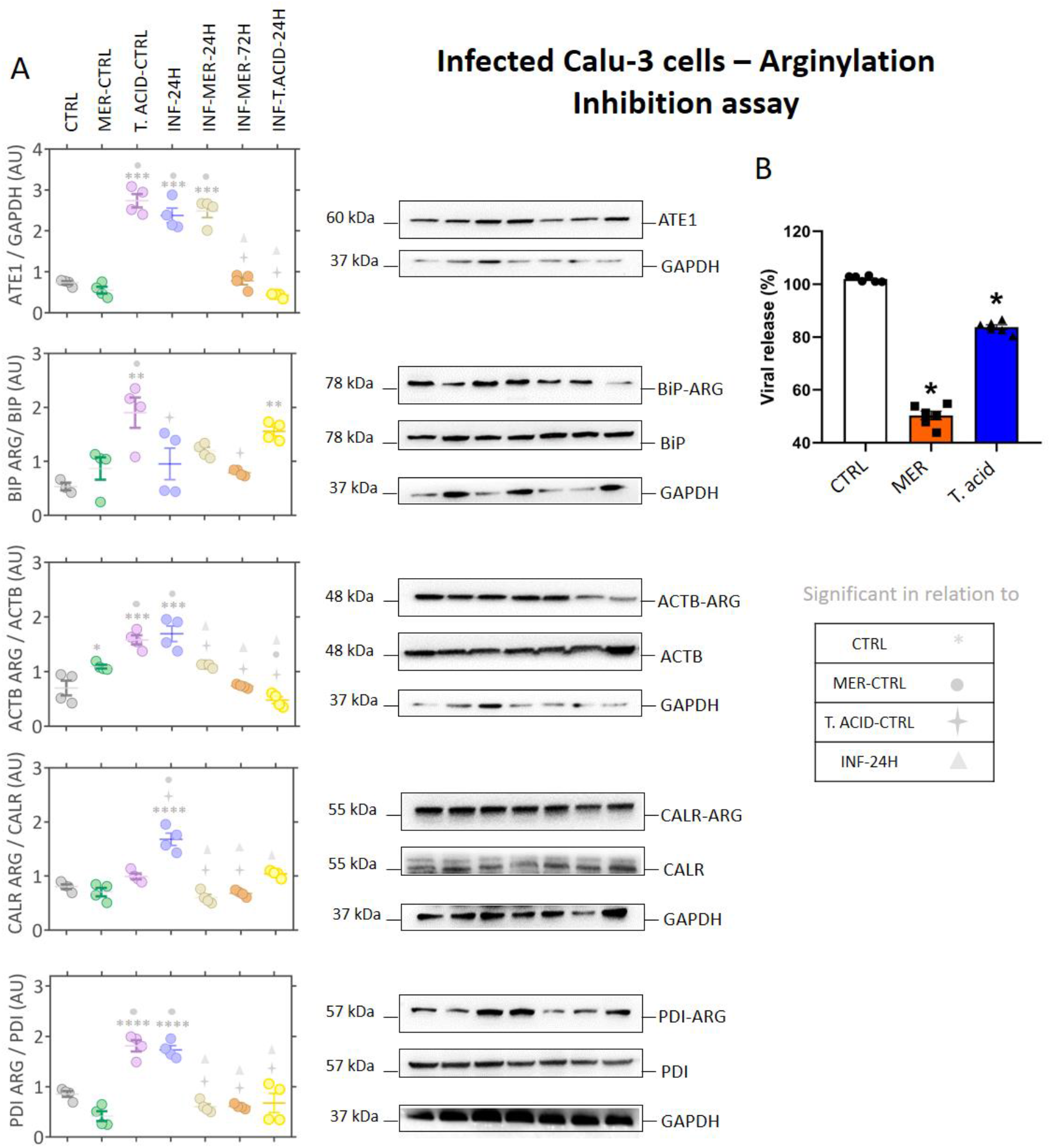
ATE1 inhibition assay in infected cells. (**A**) Representative images of western blotting analysis performed on Calu-3 cells for the proteins ATE1/GAPDH, R-BiP/BiP, R-CALR/CALR, and R-PDI/PDI; (**B**) PCR results for viral load after exposure to the inhibitors. The ATE1 inhibitors merbromin (MER, 25μM) and tannic acid (T. acid, 1μM) were evaluated. Each point represents an independent experiment (n = 3).

### Single-cell RNA-seq data showed that macrophages and epithelial cells express ATE1

After verifying the expression and subcellular location of proteins involved in the arginylation process, we investigated which cell types express ATE1. A reanalysis of single-cell RNASeq data published by Chua et. al^58^ was conducted using INF group consisted by critically ill patients, hospitalized for more than 20 days or who died due to the progression of COVID-19, and CTRL group of not infected with the virus using nasopharyngeal/pharyngeal swabs samples. A total of 17 cell clusters were identified (**Fig. 7A**). Observing the representativeness of each patient in the cell clusters, it was possible to verify that clusters 4, 7, and 8 presented a statistical difference (p < 0.05) between the INF and CTRL groups (**Fig. 7B**). We identified clusters 4, 9, 15, and 17 as clusters expressing ATE1 (**Fig. S2A**). The top five markers of cluster 4 are *LYZ, SRGN, HLA-DPB1, CD74*, and *TYROBP*, all markers of macrophages (**Supplementary File 4**). By tracking the classical macrophage markers: *MARCO*^71^, *CD163*^72^, *MRC1*^73^, and *MSR1*^74^, the presence of macrophages in cluster 4 was reinforced (**Fig. S2**). The macrophages present in cluster 4 were isolated and the genes differentially regulated between the CTRL and INF groups were determined (**Fig. S2B**). The expressions of *ATE1, CALR, ACTB, PDIA3, PDIA6*, and *PDIA4* genes did not show statistical significance between the groups (**Fig. S2C**). However, the *BiP/HSPA5* gene expression was increased in the INF group. The upregulated genes in cluster 4 were associated with interferon type I induction and signaling during SARS-CoV-2 infection, pulmonary fibrosis, proteasome degradation, and ferroptosis. On the other hand, the downregulated genes were related to peptide chain elongation, oxidative phosphorylation, and MHC class II complex (**Fig. S3**). To verify the modulation of arginylation, a western blotting was performed in macrophages infected with SARS-CoV-2 (**Fig. 7C**). We confirmed that there was no difference in the ATE1 modulation between the CTRL and INF groups after 24h and 48h of infection. In addition, the inhibitors induced an increase in ATE1 enzyme levels in macrophages at 48h (MER) and 24h (tannic acid) after treatment. The abilities of tannic acid to decrease arginylation levels occurred mainly after 48h of infection (**Fig. 7C**). These data suggested that the arginylation behavior in infected macrophages was different from that observed in Vero CCL-81 and Calu-3 cells. The expressions of ER chaperones, CALR and BIP, and ACTB showed statistical significance between the INF (48h) and CTRL groups in western blotting assays (**Fig. 7C**), with reduced expressions in INF group. Looking at differentially regulated genes between the CTRL and INF groups in clusters 9, 15, and 17 (**Fig. S4**), we identified a statistically significant increase of *ATE1* in the INF group in cluster 15, which was enriched mainly with epithelial cell markers (**Supplementary File 4**).

**Figure 7.**
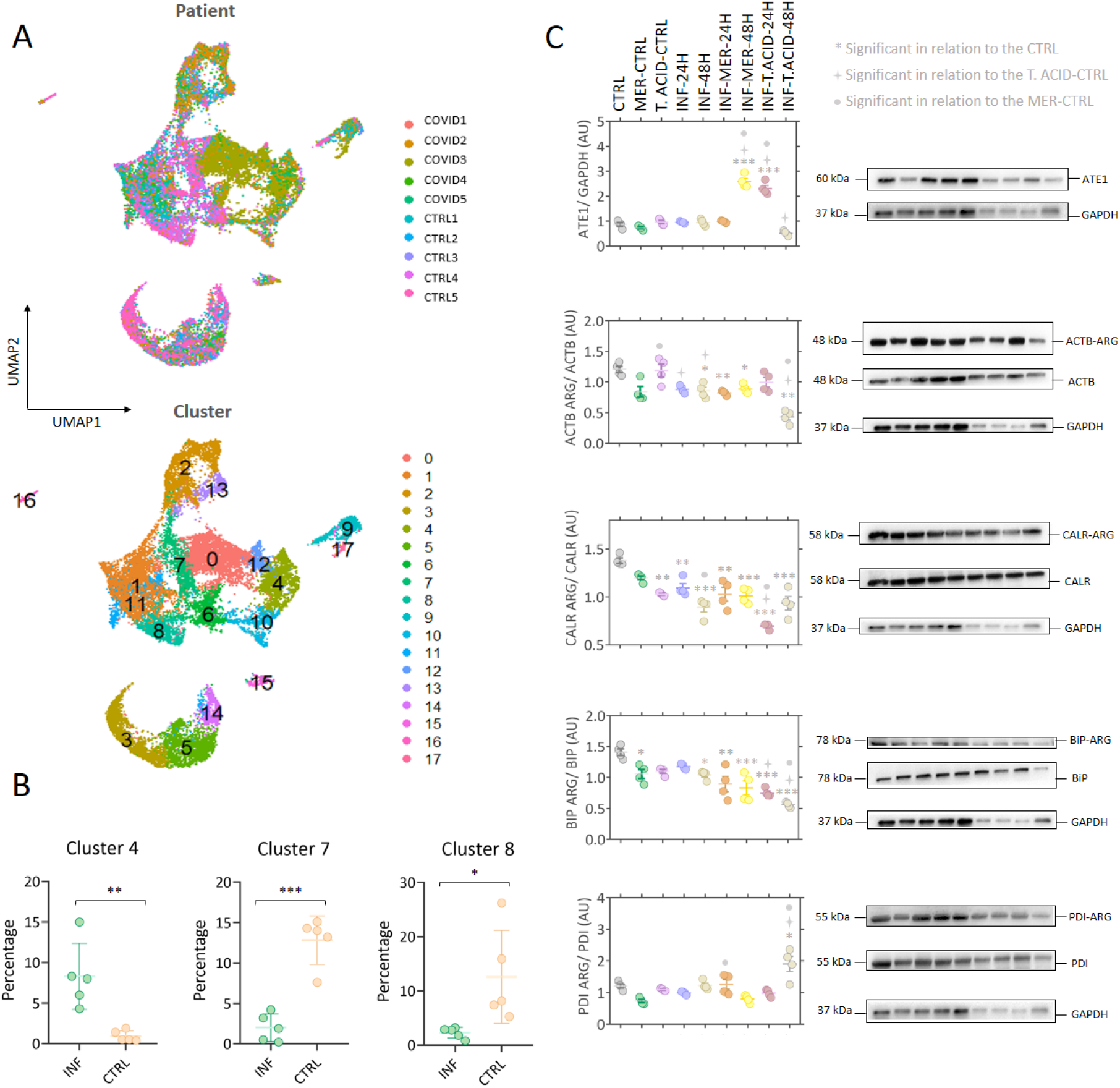
Single-cell RNA-seq analysis of nasopharyngeal samples. **(A)** Single-cell RNA-seq analysis indicating cell clustering by patients and by cell type; (**B**) Differentially regulated clusters (p-value < 0.05) between the infected (INF) and control (CTRL) groups; (**C**) Representative images of western blotting analysis of infected macrophages, indicating the modulation of ATE1 and arginylated and total CALR, PDI, BiP, and ACTB proteins.

### N-end rule pathway was regulated in SARS-CoV and MERS-CoV infections but not in H1N1 influenza and Respiratory syncytial virus (RSV) infections

We verified whether modulation of the N-end rule pathway was recurrent in other respiratory viral infections or it was a specific signature of SARS-CoV-2 (**Fig. S5**). The identification/regulation of proteins related to N-terminal methionine removal and ubiquitination was less recurrent in influenza, RSV, and human adenovirus infections. On the other hand, viruses of the *Coronaviridae* family, such as SARS-CoV and MERS-CoV, showed modulation of proteins involved in these reactions. Convergently, ATE1 was not identified or it was downregulated in *in vitro* models infected with H1N1 and RSV; in contrast, it was upregulated in cells infected with SARS-CoV and MERS-CoV. These data indicated an arginylation-dependent signature during infection with viruses from the *Coronaviridae* family.

## Discussion

In this study, we confirmed the modulation of the N-end rule pathway and proteins arginylation during SARS-CoV-2 infection by a combined *in silico* analysis of multi-omic studies and orthogonal experimental validation (**Fig.1**). We demonstrated, for the first time, an increase of ATE1expression, a critical enzyme involved in arginylation, during *in vitro* SARS-CoV-2 infection (**Fig.2**). In fact, in human Calu-3 cells, a progressive increase of ATE1 expression was observed after 6h of SARS-CoV-2 infection, when an increase of viral proteins has been previously demonstrated^46^. Interestingly, increased ATE1 expression was also observed in a monkey-derived cell line (Vero E6) indicating that this modulation may occur independent to species. Moreover, ATE1 expression increase was also demonstrated in MERS-CoV (24h) and SARS-CoV (36h) infections^51^, but not in other respiratory viruses infections such as RSV^75, 54, 76^ and influenza^49, 50^, suggesting that involvement of the N-end rule pathway may be a molecular signature specific for the *Coronaviridae* family.

A previous study by our group revealed an activation of the UPR pathway after 6h of SARS-CoV-2 infection^17^, corroborating previous observation of ER-stress enhancement^26^ and UPR pathway activation during viral infection^18, 19^. Based on these findings, we hypothesized that increased misfolded or unfolded proteins produced during SARS-CoV-2 infection may be tagged for degradation by arginylation in order to maintain cellular homeostasis. In fact, our *in silico* multiomic analysis identified several ER-related proteins associated with ATE1 expression (**Fig.3**), including BiP/HSPA5, PDI and CALR, and we followed their arginylation rate along with the infection (**Fig.4**). As expected, BiP/HSPA5 expression increased 48hs after infection in both human and monkey cell lines. It has been shown that the viral spike glycoprotein (S) plays a fundamental role in SARS-CoV-2 infection in the process of receptor recognition and cell membrane fusion^2^, and it induced the transcriptional activation of Hsp90β member 1 and BiP/HSPA5 chaperones^14^. The increased expression of these chaperones has resulted in increased folding and processing of abundantly expressed proteins during SARS-CoV replication^20, 77^. Moreover, BiP/HSPA5 arginylation has also been induced by transient transfection of several dsDNAs, suggesting that its modulation may be also related to the detection of pathogenic dsDNA and activation of the immune system^78^. On the other hand, we observed a reduction in the levels of CALR, a protein involved in the folding and maturation of glycoproteins^79, 80^. CALR decrease has also been described in other viral infections (influenza virus, SFV, or VSV) leading to accelerate maturation of cellular and viral glycoproteins, with a modest decrease in the folding efficiency^81^. We speculated that the progressive reduction in CALR levels in Calu-3 cells might be associated with acceleration of the coronavirus S-glycoprotein maturation. The receptor-binding domain of the S-glycoprotein and ACE2 has several cysteine residues. The reduction of disulfide bonds to thiol decreased the binding affinity interaction^82^. The PDI protein is a redox-regulated chaperone related to the formation, isomerization, and reduction of disulfide bonds^83^. Our multi-omic analysis identified the downregulation of PDI members and immunoblot data points to a significant reduction in PDI levels 6h after infection, suggesting that the interaction between viral S-glycoprotein and ACE2 is not being affected by the dependent redox status of PDI. The UPR pathway is activated to restore ER homeostasis, and in case of failure, apoptotic events are induced^84^. The virus can also hijack the host’s ubiquitination machinery^85^; however, the biological functions of this action are still unknown.

Our multi-omic analysis demonstrated that the arginylation-related proteins were mainly located in the ER, convergent to our *in vitro* results of the proteins involved in the UPR pathway. Additionally, the arginylation-related proteins were also located in the cytoskeleton, a key structure in the host-pathogen interaction^86^ (**Fig. 5**). Cytoskeleton proteins participate throughout the viral replication cycle, as SARS-CoV-2 enters into target cells using intermediate filament proteins, sequesters microtubules to transport itself to replication/assembly sites, and promotes polymerization of actin filaments to exit the cell^87, 88^. Moreover, cytoskeleton proteins were among the experimentally arginylated proteins identified in previous studies (Seo et al.^56^ and Wong et al.^57^). We monitored the arginylation levels of one major cytoskeleton protein, ACTB, and we observed its increase in the first 2h after infection in Calu-3 cells, with a decrease at 48h, when apoptosis-related proteins were activated^17^. The ACTB arginylation pattern was different in infected Vero cells, as was already observed for phosphorylation^89^. Such observed difference stressed the importance to use multiple cell models to assess the cellular consequences of post-translational modifications in SARS-CoV-2 infection.

Our analysis of single-cell RNA-seq data revealed a cellular compartment consisted by macrophage in infected patients. In fact, the characterization of immune cells in bronchoalveolar lavage fluid has shown that pro-inflammatory monocyte-derived macrophages were more abundant in patients with severe SARS-Cov-2 infection than in those with moderate disease or healthy individuals. Furthermore, critically ill patients presented a lower proportion of myeloid dendritic cells, plasmacytoid dendritic cells, and T-cells than patients with mild infection^90^. We also demonstrated ATE1 protein expression in macrophages. However, no difference in the abundance of proteins related to arginylation was detected comparing CTRL and INF groups (**Fig 7**). In fact, previous study has shown that SARS-CoV-2 was capable of infecting macrophages without causing any cytopathic effect, and the virus was also capable of inducing host immunoparalysis^91^. Moreover, we also identified a cellular compartment with epithelial cell markers presenting significant increase of *ATE1* expression, consistent with previous observation of ATE1 protein expression in the main lung epithelial cells, ciliary cells type 1 and 2, infected by SARS-CoV-2^92^. ATE1 expression in lung epithelial cells was higher in SARS-CoV-2 infected patients compared to controls.

Notably, we found that arginylation inhibitors, tannic acid and MER decreased viral load or prevented viral entry into the cell. Furthermore, our assays indicated a decrease in the abundance of ATE1 (**Fig.6**). Tannic acid was recently described as a potent inhibitor of SARS-CoV-2, through the thermodynamically stable binding with the proteins Mpro and TMPRSS2, crucial for the entry of the virus into the cell^93^. Here, we confirmed the potential of tannic acid to reduce viral load, and furthermore, to modulate ATE1 levels during infection. In addition, we also demonstrated for the first time that, like tannic acid, the arginylation inhibitor MER was also capable of reducing both viral load and ATE1 level. Our main findings are summarized in **Fig. 8**.

**Figure 8.**
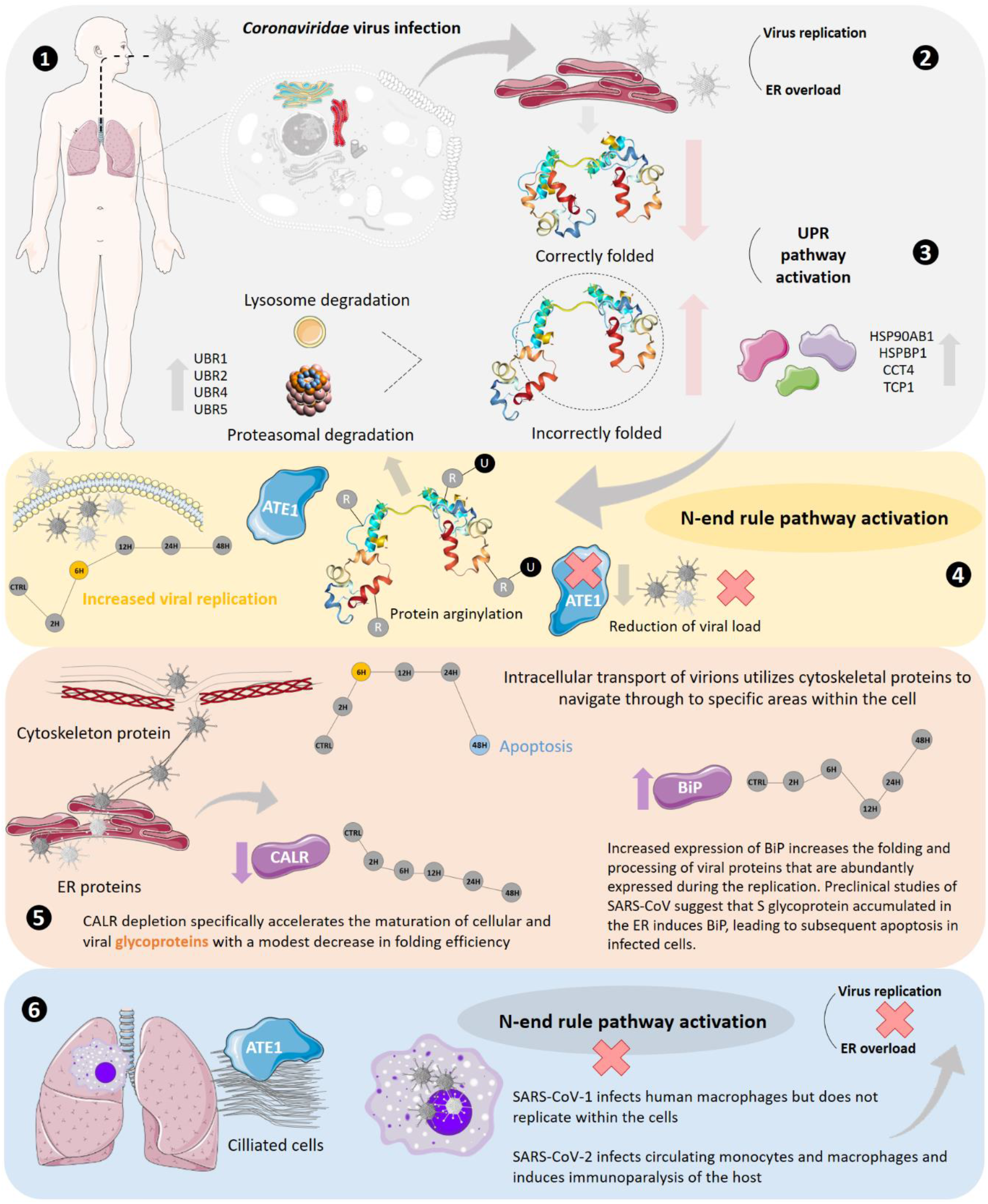
Summary of the main findings, indicating the hypothesis and results found.

## Conclusion

We report for the first time the role of protein arginylation and modulation of the N-end rule pathway during SARS-CoV-2 infection. Differential regulation of proteins involved in all reactions that make up the N-end rule pathway was demonstrated, with emphasis on the upregulation of the ATE1 protein, evidenced by omic data and western blotting. Furthermore, we show that proteins that have their levels correlated with ATE1 perform biological functions related to chaperone activity and binding to unfolded proteins. An important finding revealed that modulation of the N-end rule pathway differs between different types of infected cells, such as macrophages, Vero CCL-81, and Calu-3 cells. Finally, we show the importance of this process through reducing viral load using tannic acid and MER, known arginylation inhibitors.

## Acknowledgments

Prof. Francisco Laurindo from Faculdade de Medicina USP is acknowledged for providing the PDI antibody. We are grateful for the financial support provided by the São Paulo Research Foundation (FAPESP, grants processes n° 2018/18257-1 (GP), 2018/15549-1 (GP), 2021/00140-3 (JMDS), 2020/02988-7 (SKNM); 2015/26722-8 (CW), 2020/12277-0 (EES), 2020/06409-1 (ELD), 2021/00507-4 (VdMG); PIBIC (Programa Institucional de Bolsas de Iniciação Científica) 1750 to CMSM, by the Conselho Nacional de Desenvolvimento Científico e Tecnológico, 140428/2021-6 to BRB, 870219/1997-9 to VFS, (“Bolsa de Produtividade” (SKNM and 307854/2018-3 GP, 302917/2019-5 CRFM)); by FMUSP (SKNM); by the Coordenação de Aperfeiçoamento de Pessoal de Nível Superior (CAPES bolsa PNPD 88887372048/2019-00 to LRF), and Brazilian Ministry of Science, Technology and Innovation (MCTI-Rede Virus).

## Contributions

JMdS, LRF, VdMG, VFS, CMSM, BRB, EEdS. performed the experiments. JMdS. GP. designed the research. JMdS, LRF, VdMG, BRB, EEdS conducted and analysed SARS-CoV-2 infections. JMdS, CMSM and VFS performed western blotting analysis. ELD, CRF, CW supervised the viral isolation, titration and infection. JMdS, SKNM and GP wrote the first version of the manuscript. All authors have critically reviewed the manuscript. JMdS and GP conceived and directed the project.

## Competing interests

The authors declare no competing interests.

## Supplementary Figures

**Figure S1.**
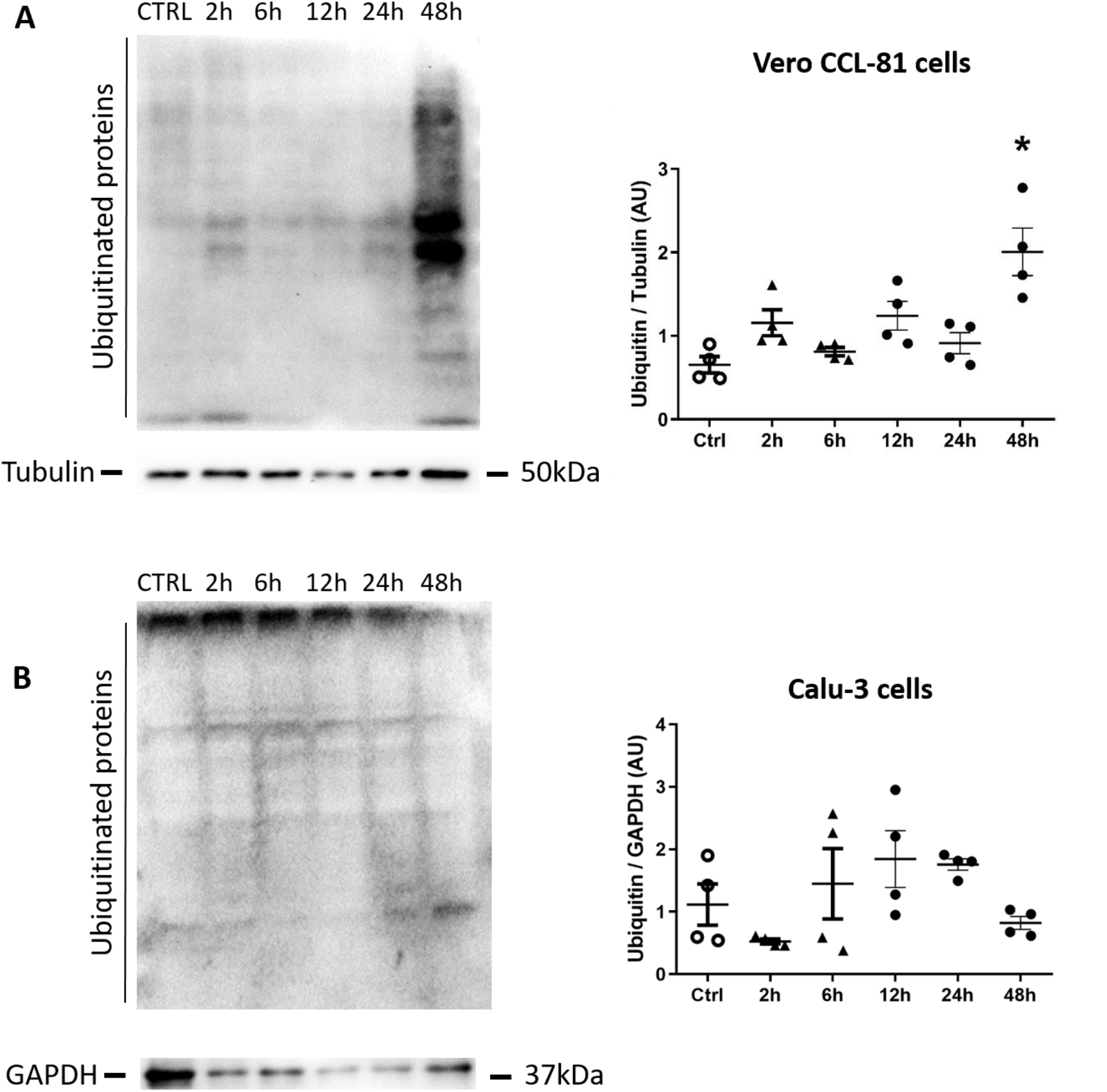
Representative images of western blotting performed to verify ubiquitination in Vero CCL-81 (**A**) and Calu-3 (**B**) cells.

**Figure S2.**
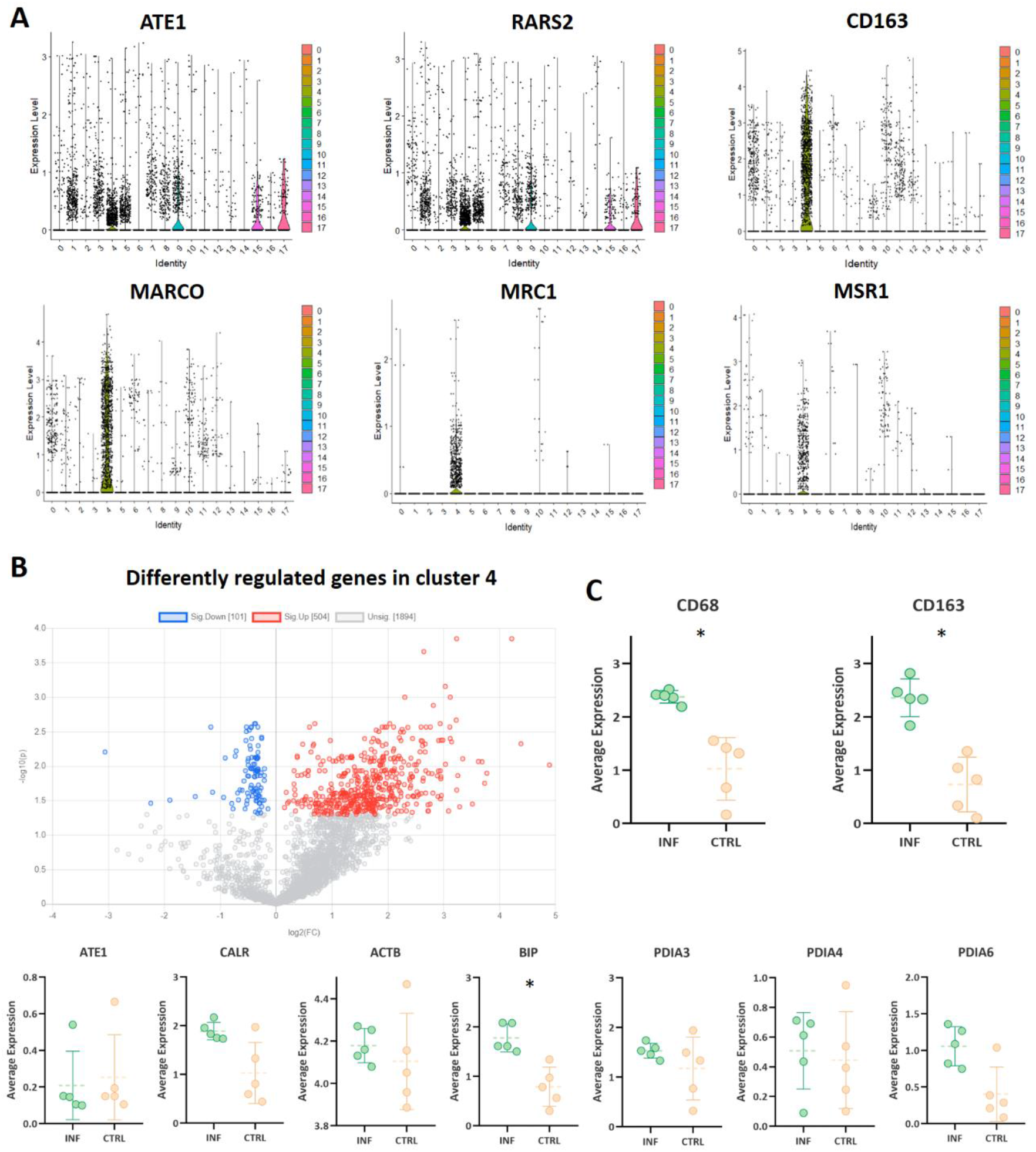
Expression of *ATE1, RARS2, CD163, MARCO, MRC1*, and *MSR1* in cell clusters identified by reanalysis of single-cell RNA-seq data (**A**); Volcano plot showing the differentially regulated genes in cluster 4 (INF/CTRL). Red and blue colors show up-regulated and down-regulated genes, respectively (q-value <0.05). The gray color indicates genes that did not show statistical significance (**B**); Regulation of genes *CD68, CD163, ATE1, CALR, PDI, BiP, PDIA3, PDIA4*, and *PDIA6* in cluster 4 (C). The (*) symbol indicates that the gene is differently regulated.

**Figure S3.**
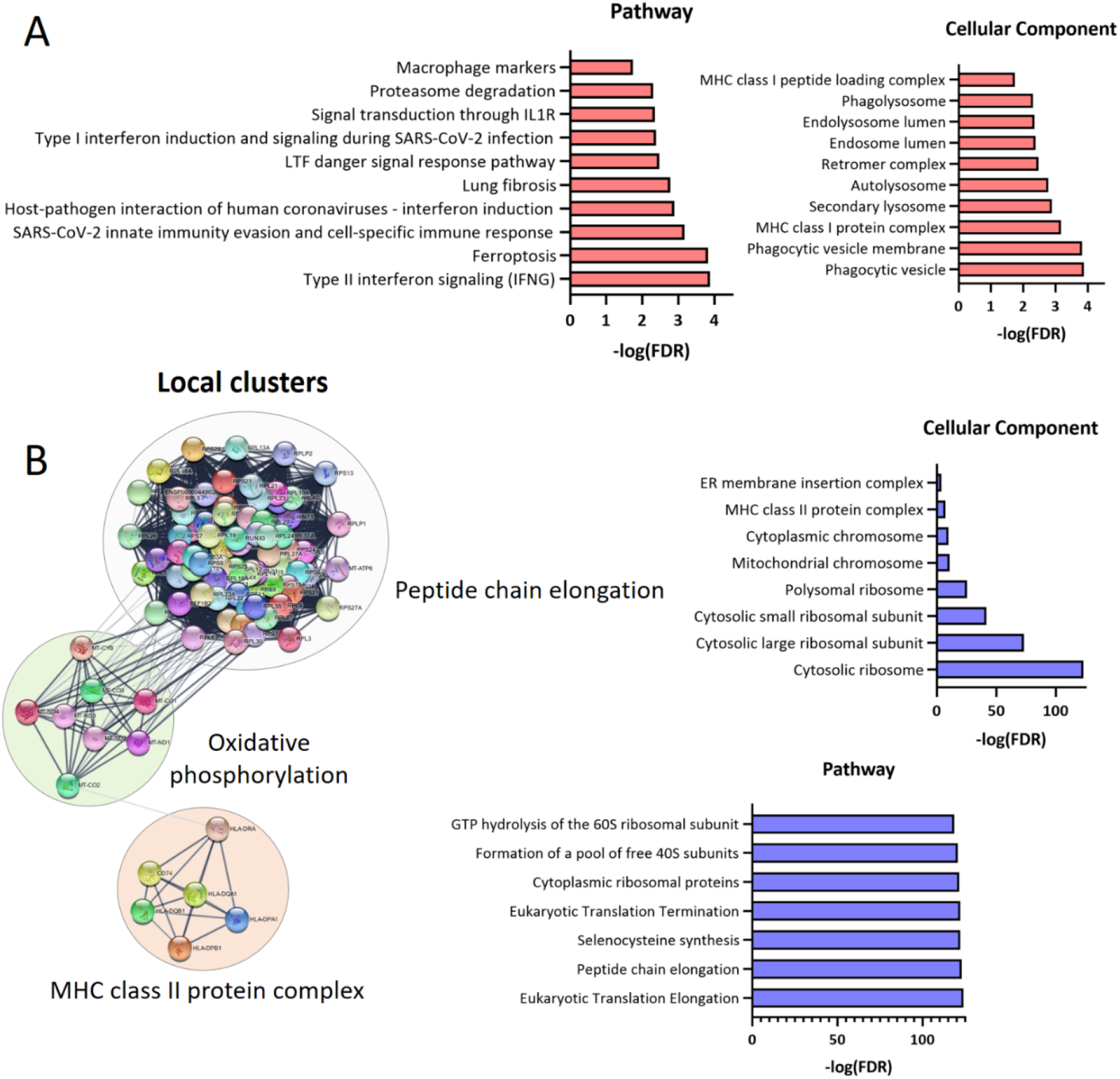
Pathways and cellular components related to upregulated genes in cluster 4 (**A**); Downregulated genes connectivity according to biological process and enrichment by cellular components and pathways (**B**). All listed ontologies have statistical significance (q-value < 0.01 - Benjamini-Hochberg).

**Figure S4.**
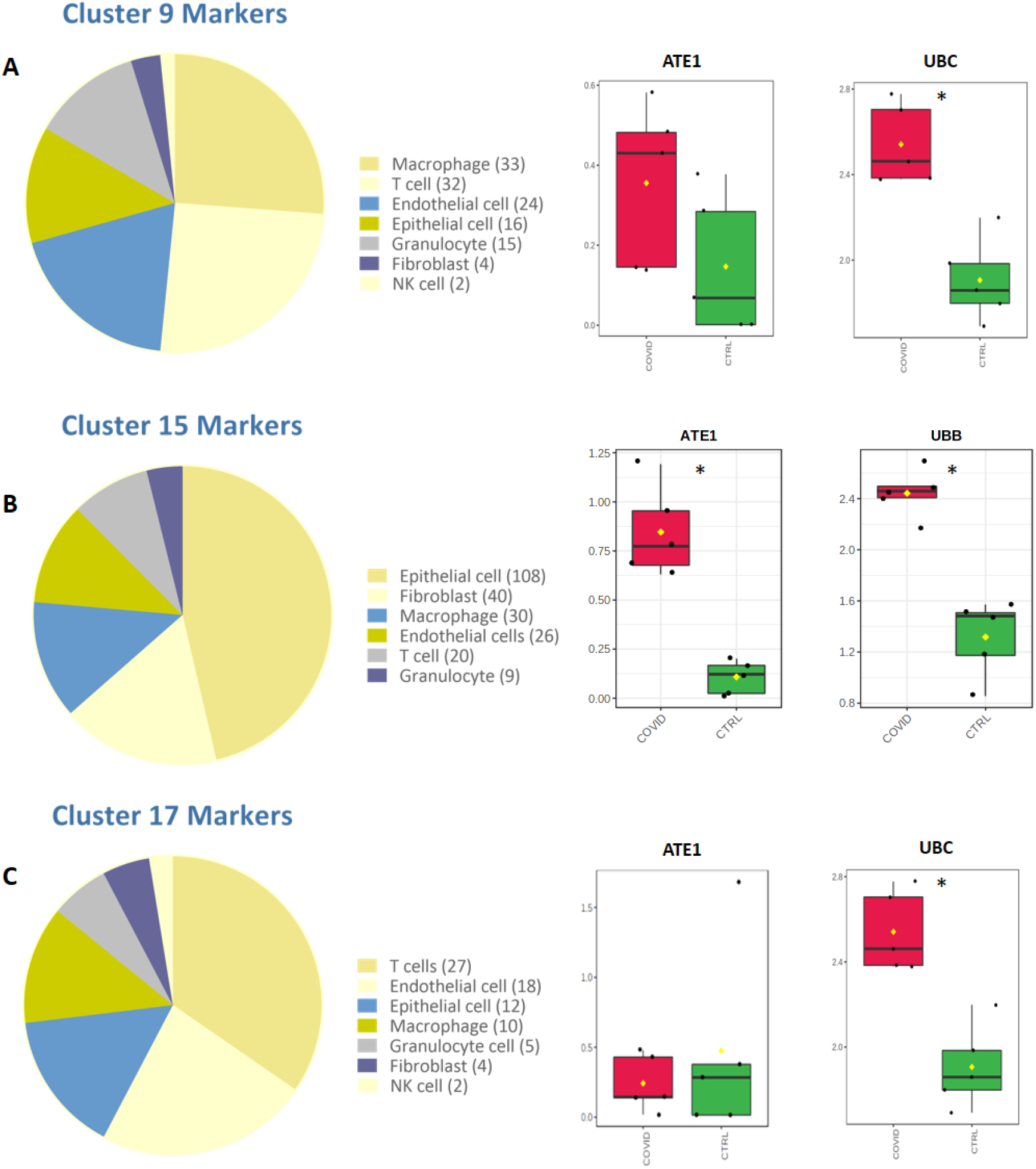
Cell markers, *ATE1* and polyubiquitins *UBB* or *UBC* expressions in cluster 9 (**A**), cluster 15 (**B**), and cluster (**17**). The (*) symbol indicates that the gene is differentially regulated.

**Figure S5.**
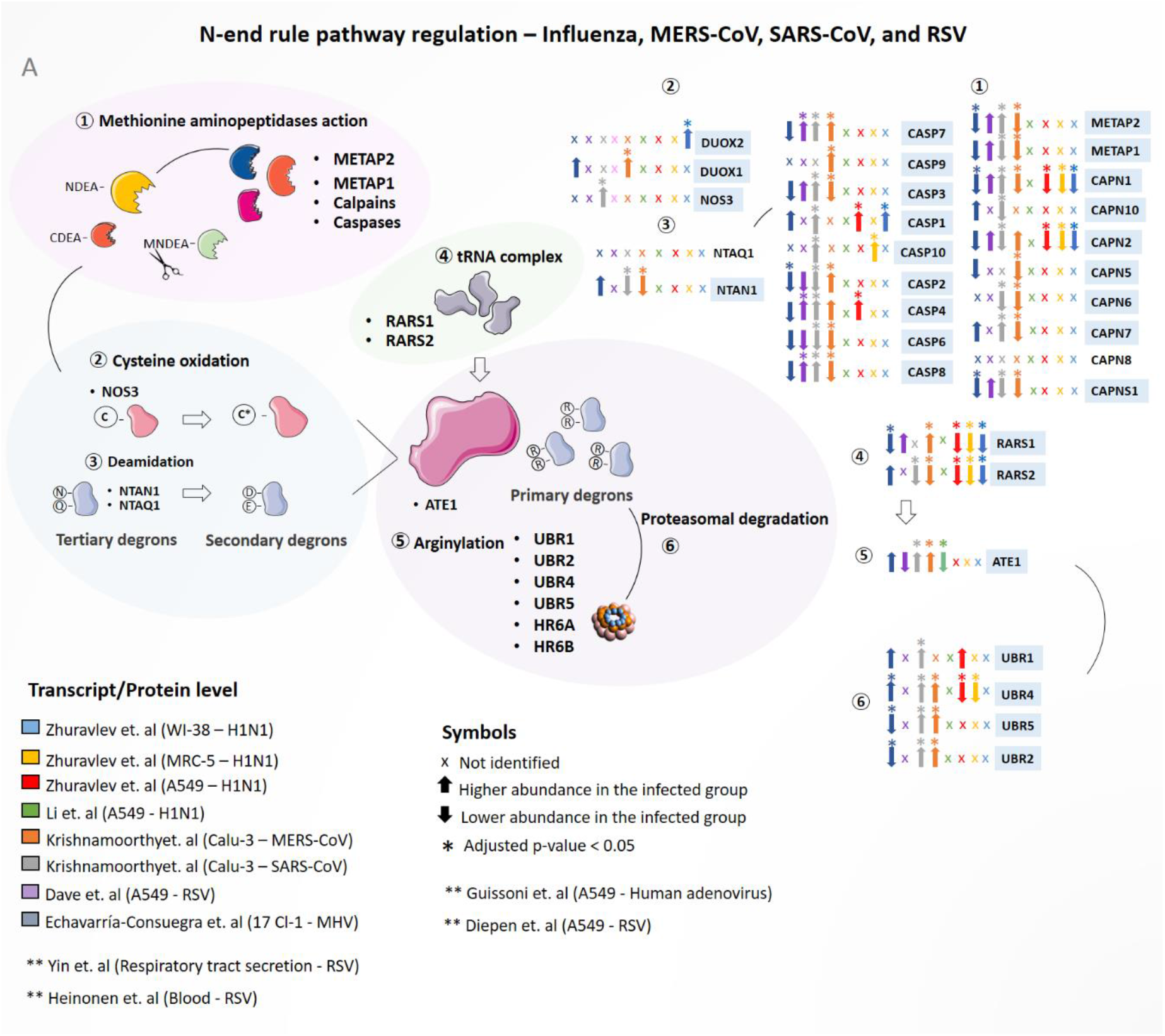
Modulation of proteins involved in the N-end rule pathway during H1N1 influenza virus, A549, MERS-CoV, and SARS-CoV. Proteins were considered differentially regulated if they had a q-value < 0.05 (Benjamini-Hochberg). Up arrows indicate proteins with the higher abundance in the infected group (INF), and down arrows indicate proteins with lower abundance in the control group (CTRL).

## Supplementary Files

**Supplementary File 1**. N-end rule pathway (Zecha et. al 2020) (A); Potential to be arginylated (Human proteome) (B); Potential to be arginylated (Zecha et. al 2020) (C); Differentially regulated proteins (Saccon et. al) (D); Differentially regulated proteins (Nie et. al) (E); Differentially regulated proteins (Leng et. al) (F); Differentially regulated proteins (Qiu et. al) (G); Differentially regulated proteins (Bojkova et. al) (H); Differently regulated transcripts (Wu et. al) (I) and Differentially regulated transcripts (Desai et. al) (J).

**Supplementary File 2**. Proteins and genes correlated with ATE1 in Saccon et. al (A); Nie et. al (B); Qiu et. al (C); Bojkova et. al (D); Wu et. al (E); and Desai et. al (F); Venn diagram of correlated proteins (G); and Gene ontology analysis of correlated proteins (H).

**Supplementary File 3**. Subcellular location of proteins correlated with ATE1 (At least 2 studies) (A); Subcellular location of proteins correlated with ATE1 (At least 3 studies) (B); Differentially regulated arginylated proteins (experimentally validated) (C) and List of 152 arginylated proteins experimentally validated.

**Supplementary File 4**. All markers for each clusters (A); matrix expression for: cluster 4 (B); cluster 9 (C); cluster 15 (D); and cluster 17 (E); Volcano plot results for: cluster 4 (F); cluster 9 (G); cluster 15 (H); and cluster 17 (I); Cell markers for: cluster 9 (J); cluster 15 (K); and cluster 17 (L); Gene ontology analysis of upregulated (M) and downregulated (N) genes in cluster 4.

## Notes

### Competing Interest Statement

The authors have declared no competing interest.

## References

1. Mahalmani, V. M. et al. COVID-19 pandemic: A review based on current evidence. Indian J Pharmacol 52, 117–129 (2020).

2. Huang, C. et al. Clinical features of patients infected with 2019 novel coronavirus in Wuhan, China. The Lancet 395, 497–506 (2020).

3. Holshue, M. L. et al. First Case of 2019 Novel Coronavirus in the United States. N Engl J Med 382, 929–936 (2020).

4. Gandhi, R. T., Lynch, J. B. & del Rio, C. Mild or Moderate Covid-19. N Engl J Med 383, 1757–1766 (2020).

5. Esakandari, H. et al. A comprehensive review of COVID-19 characteristics. Biol Proced Online 22, 19 (2020).

6. He, X. et al. Clinical Symptom Differences Between Mild and Severe COVID-19 Patients in China: A Meta-Analysis. Front. Public Health 8, 561264 (2021).

7. Yuen, K.-S., Ye, Z.-W., Fung, S.-Y., Chan, C.-P. & Jin, D.-Y. SARS-CoV-2 and COVID-19: The most important research questions. Cell Biosci 10, 40 (2020).

8. Petrosillo, N., Viceconte, G., Ergonul, O., Ippolito, G. & Petersen, E. COVID-19, SARS and MERS: are they closely related? Clin Microbiol Infect 26, 729–734 (2020).

9. Guo, Q. & He, Z. Prediction of the confirmed cases and deaths of global COVID-19 using artificial intelligence. Environ Sci Pollut Res Int 28, 11672–11682 (2021).

10. Harrison, A. G., Lin, T. & Wang, P. Mechanisms of SARS-CoV-2 Transmission and Pathogenesis. Trends Immunol 41, 1100–1115 (2020).

11. Rabaan, A. A. et al. SARS-CoV-2, SARS-CoV, and MERS-COV: A comparative overview. Infez Med 28, 174–184 (2020).

12. Sixto-López, Y. et al. Structural insights into SARS-CoV-2 spike protein and its natural mutants found in Mexican population. Sci Rep 11, 4659 (2021).

13. Hatmal, M. M. et al. Comprehensive Structural and Molecular Comparison of Spike Proteins of SARS-CoV-2, SARS-CoV and MERS-CoV, and Their Interactions with ACE2. Cells 9, E2638 (2020).

14. Chan, C.-P. et al. Modulation of the unfolded protein response by the severe acute respiratory syndrome coronavirus spike protein. J Virol 80, 9279–9287 (2006).

15. Fung, T. S., Huang, M. & Liu, D. X. Coronavirus-induced ER stress response and its involvement in regulation of coronavirus-host interactions. Virus Res 194, 110–123 (2014).

16. Köseler, A., Sabirli, R., Gören, T., Türkçüer, İb. & Kurt, Ö. Endoplasmic Reticulum Stress Markers in SARS-COV-2 Infection and Pneumonia: Case-Control Study. In Vivo 34, 1645–1650 (2020).

17. Rosa-Fernandes, L. et al. SARS-CoV-2 activates ER stress and Unfolded protein response. http://biorxiv.org/lookup/doi/10.1101/2021.06.21.449284 (2021) doi:10.1101/2021.06.21.449284.

18. Rashid, F., Dzakah, E. E., Wang, H. & Tang, S. The ORF8 protein of SARS-CoV-2 induced endoplasmic reticulum stress and mediated immune evasion by antagonizing production of interferon beta. Virus Res 296, 198350 (2021).

19. Balakrishnan, B. & Lai, K. Modulation of SARS-CoV-2 Spike-induced Unfolded Protein Response (UPR) in HEK293T cells by selected small chemical molecules. http://biorxiv.org/lookup/doi/10.1101/2021.02.04.429769 (2021) doi:10.1101/2021.02.04.429769.

20. Aoe, T. Pathological Aspects of COVID-19 as a Conformational Disease and the Use of Pharmacological Chaperones as a Potential Therapeutic Strategy. Front. Pharmacol. 11, 1095 (2020).

21. Barabutis, N. Unfolded Protein Response in Acute Respiratory Distress Syndrome. Lung 197, 827–828 (2019).

22. Bouchecareilh, M. & Balch, W. E. Proteostasis: a new therapeutic paradigm for pulmonary disease. Proc Am Thorac Soc 8, 189–195 (2011).

23. Lecker, S. H., Goldberg, A. L. & Mitch, W. E. Protein degradation by the ubiquitin-proteasome pathway in normal and disease states. J Am Soc Nephrol 17, 1807–1819 (2006).

24. Wang, X. & Robbins, J. Proteasomal and lysosomal protein degradation and heart disease. J Mol Cell Cardiol 71, 16–24 (2014).

25. Ravid, T. & Hochstrasser, M. Diversity of degradation signals in the ubiquitin-proteasome system. Nat Rev Mol Cell Biol 9, 679–690 (2008).

26. Livneh, I., Kravtsova-Ivantsiv, Y., Braten, O., Kwon, Y. T. & Ciechanover, A. Monoubiquitination joins polyubiquitination as an esteemed proteasomal targeting signal. Bioessays 39, (2017).

27. Tasaki, T., Sriram, S. M., Park, K. S. & Kwon, Y. T. The N-end rule pathway. Annu Rev Biochem 81, 261–289 (2012).

28. Varshavsky, A. The N-end rule pathway and regulation by proteolysis. Protein Sci 20, 1298–1345 (2011).

29. Kopitz, J., Rist, B. & Bohley, P. Post-translational arginylation of ornithine decarboxylase from rat hepatocytes. Biochem J 267, 343–348 (1990).

30. Eriste, E. et al. A novel form of neurotensin post-translationally modified by arginylation. J Biol Chem 280, 35089–35097 (2005).

31. Soffer, R. L. Enzymatic arginylation of beta-melanocyte-stimulating hormone and of angiotensin II. J Biol Chem 250, 2626–2629 (1975).

32. Wang, J. et al. Protein arginylation targets alpha synuclein, facilitates normal brain health, and prevents neurodegeneration. Sci Rep 7, 11323 (2017).

33. Hu, R.-G., Wang, H., Xia, Z. & Varshavsky, A. The N-end rule pathway is a sensor of heme. Proceedings of the National Academy of Sciences 105, 76–81 (2008).

34. Cha-Molstad, H., Kwon, Y. T. & Kim, B. Y. Amino-terminal arginylation as a degradation signal for selective autophagy. BMB Rep 48, 487–488 (2015).

35. Varshavsky, A. N-degron and C-degron pathways of protein degradation. Proc Natl Acad Sci USA 116, 358–366 (2019).

36. Wang, J. et al. Arginyltransferase ATE1 catalyzes midchain arginylation of proteins at side chain carboxylates in vivo. Chem Biol 21, 331–337 (2014).

37. Wang, J. et al. Target site specificity and in vivo complexity of the mammalian arginylome. Sci Rep 8, 16177 (2018).

38. Deka, K. & Saha, S. Arginylation: a new regulator of mRNA stability and heat stress response. Cell Death Dis 8, e2604 (2017).

39. Kumar, A. et al. Posttranslational arginylation enzyme Ate1 affects DNA mutagenesis by regulating stress response. Cell Death Dis 7, e2378–e2378 (2016).

40. Deka, K., Singh, A., Chakraborty, S., Mukhopadhyay, R. & Saha, S. Protein arginylation regulates cellular stress response by stabilizing HSP70 and HSP40 transcripts. Cell Death Discovery 2, 16074 (2016).

41. Saha, S. & Kashina, A. Posttranslational arginylation as a global biological regulator. Dev Biol 358, 1–8 (2011).

42. Saccon, E. et al. Cell-type-resolved quantitative proteomics map of interferon response against SARS-CoV-2. iScience 24, 102420 (2021).

43. Nie, X. et al. Multi-organ proteomic landscape of COVID-19 autopsies. Cell 184, 775–791.e14 (2021).

44. Leng, L. et al. Pathological features of COVID-19-associated lung injury: a preliminary proteomics report based on clinical samples. Signal Transduct Target Ther 5, 240 (2020).

45. Qiu, Y. et al. Postmortem Tissue Proteomics Reveals the Pathogenesis of Multiorgan Injuries of COVID-19. https://www.researchsquare.com/article/rs-38091/v1 (2020) doi:10.21203/rs.3.rs-38091/v1.

46. Bojkova, D. et al. Proteomics of SARS-CoV-2-infected host cells reveals therapy targets. Nature 583, 469–472 (2020).

47. Wu, M. et al. Transcriptional and proteomic insights into the host response in fatal COVID-19 cases. Proc Natl Acad Sci U S A 117, 28336–28343 (2020).

48. Desai, N. et al. Temporal and spatial heterogeneity of host response to SARS-CoV-2 pulmonary infection. Nat Commun 11, 6319 (2020).

49. Zhuravlev, E. et al. RNA-Seq transcriptome data of human cells infected with influenza A/Puerto Rico/8/1934 (H1N1) virus. Data Brief 33, 106604 (2020).

50. Li, J. et al. Transcriptome Profiling Reveals Differential Effect of Interleukin-17A Upon Influenza Virus Infection in Human Cells. Front Microbiol 10, 2344 (2019).

51. Krishnamoorthy, P., Raj, A. S., Roy, S., Kumar, N. S. & Kumar, H. Comparative transcriptome analysis of SARS-CoV, MERS-CoV, and SARS-CoV-2 to identify potential pathways for drug repurposing. Comput Biol Med 128, 104123 (2021).

52. Ampuero, S. et al. Time-course of transcriptome response to respiratory syncytial virus infection in lung epithelium cells. Acta Virol 62, 310–325 (2018).

53. Besteman, S. B. et al. Transcriptome of airway neutrophils reveals an interferon response in life-threatening respiratory syncytial virus infection. Clinical Immunology 220, 108593 (2020).

54. Dave, K. A. et al. A comprehensive proteomic view of responses of A549 type II alveolar epithelial cells to human respiratory syncytial virus infection. Mol Cell Proteomics 13, 3250–3269 (2014).

55. Zecha, J. et al. Data, Reagents, Assays and Merits of Proteomics for SARS-CoV-2 Research and Testing. Mol Cell Proteomics 19, 1503–1522 (2020).

56. Seo, T. et al. R-catcher, a potent molecular tool to unveil the arginylome. Cell. Mol. Life Sci. 78, 3725–3741 (2021).

57. Wong, C. C. L. et al. Global analysis of posttranslational protein arginylation. PLoS Biol 5, e258 (2007).

58. Chua, R. L. et al. COVID-19 severity correlates with airway epithelium–immune cell interactions identified by single-cell analysis. Nat Biotechnol 38, 970–979 (2020).

59. Wickham, H. et al. Welcome to the Tidyverse. JOSS 4, 1686 (2019).

60. Charif, D. & Lobry, J. R. SeqinR 1.0-2: A Contributed Package to the R Project for Statistical Computing Devoted to Biological Sequences Retrieval and Analysis. in Structural Approaches to Sequence Evolution (eds. Bastolla, U., Porto, M., Roman, H. E. & Vendruscolo, M.) 207–232 (Springer Berlin Heidelberg, 2007). doi:10.1007/978-3-540-35306-5_10.

61. Breckels, L. M., Mulvey, C. M., Lilley, K. S. & Gatto, L. A Bioconductor workflow for processing and analysing spatial proteomics data. F1000Res 5, 2926 (2016).

62. Reimand, J., Kull, M., Peterson, H., Hansen, J. & Vilo, J. g:Profiler--a web-based toolset for functional profiling of gene lists from large-scale experiments. Nucleic Acids Res 35, W193–200 (2007).

63. Huang, D. W., Sherman, B. T. & Lempicki, R. A. Systematic and integrative analysis of large gene lists using DAVID bioinformatics resources. Nat Protoc 4, 44–57 (2009).

64. Raudvere, U. et al. g:Profiler: a web server for functional enrichment analysis and conversions of gene lists (2019 update). Nucleic Acids Research 47, W191–W198 (2019).

65. Heberle, H., Meirelles, G. V., da Silva, F. R., Telles, G. P. & Minghim, R. InteractiVenn: a web-based tool for the analysis of sets through Venn diagrams. BMC Bioinformatics 16, 169 (2015).

66. Stuart, T. et al. Comprehensive Integration of Single-Cell Data. Cell 177, 1888–1902.e21 (2019).

67. Pang, Z. et al. MetaboAnalyst 5.0: narrowing the gap between raw spectra and functional insights. Nucleic Acids Research 49, W388–W396 (2021).

68. Gatto, F. et al. PMA-Induced THP-1 Macrophage Differentiation is Not Impaired by Citrate-Coated Platinum Nanoparticles. Nanomaterials (Basel) 7, E332 (2017).

69. Araujo, D. B. et al. SARS-CoV-2 isolation from the first reported patients in Brazil and establishment of a coordinated task network. Mem. Inst. Oswaldo Cruz 115, e200342 (2020).

70. Corman, V. M. et al. Detection of 2019 novel coronavirus (2019-nCoV) by real-time RT-PCR. Euro Surveill 25, (2020).

71. Palecanda, A. et al. Role of the scavenger receptor MARCO in alveolar macrophage binding of unopsonized environmental particles. J Exp Med 189, 1497–1506 (1999).

72. Lau, S. K., Chu, P. G. & Weiss, L. M. CD163: a specific marker of macrophages in paraffin-embedded tissue samples. Am J Clin Pathol 122, 794–801 (2004).

73. Xu, Z.-J. et al. The M2 macrophage marker CD206: a novel prognostic indicator for acute myeloid leukemia. Oncoimmunology 9, 1683347 (2020).

74. Guo, M. et al. Triggering MSR1 promotes JNK-mediated inflammation in IL-4-activated macrophages. EMBO J 38, (2019).

75. Yin, G.-Q. et al. Differential proteomic analysis of children infected with respiratory syncytial virus. Braz J Med Biol Res 54, e9850 (2021).

76. Heinonen, S. et al. Immune profiles provide insights into respiratory syncytial virus disease severity in young children. Sci. Transl. Med. 12, eaaw0268 (2020).

77. Paladino, L. et al. The Role of Molecular Chaperones in Virus Infection and Implications for Understanding and Treating COVID-19. JCM 9, 3518 (2020).

78. Cha-Molstad, H. et al. Amino-terminal arginylation targets endoplasmic reticulum chaperone BiP for autophagy through p62 binding. Nat Cell Biol 17, 917–929 (2015).

79. Nauseef, W. M., McCormick, S. J. & Clark, R. A. Calreticulin functions as a molecular chaperone in the biosynthesis of myeloperoxidase. J Biol Chem 270, 4741–4747 (1995).

80. Salati, S. et al. Calreticulin Ins5 and Del52 mutations impair unfolded protein and oxidative stress responses in K562 cells expressing CALR mutants. Sci Rep 9, 10558 (2019).

81. Molinari, M. et al. Contrasting functions of calreticulin and calnexin in glycoprotein folding and ER quality control. Mol Cell 13, 125–135 (2004).

82. Hati, S. & Bhattacharyya, S. Impact of Thiol–Disulfide Balance on the Binding of Covid-19 Spike Protein with Angiotensin-Converting Enzyme 2 Receptor. ACS Omega 5, 16292–16298 (2020).

83. Fraternale, A. et al. Intracellular Redox-Modulated Pathways as Targets for Effective Approaches in the Treatment of Viral Infection. IJMS 22, 3603 (2021).

84. Choi, J.-A. & Song, C.-H. Insights Into the Role of Endoplasmic Reticulum Stress in Infectious Diseases. Front. Immunol. 10, 3147 (2020).

85. Cao, Z. et al. Ubiquitination of SARS-CoV-2 ORF7a promotes antagonism of interferon response. Cell Mol Immunol 18, 746–748 (2021).

86. Haglund, C. M. & Welch, M. D. Pathogens and polymers: microbe-host interactions illuminate the cytoskeleton. J Cell Biol 195, 7–17 (2011).

87. Wen, Z., Zhang, Y., Lin, Z., Shi, K. & Jiu, Y. Cytoskeleton—a crucial key in host cell for coronavirus infection. Journal of Molecular Cell Biology 12, 968–979 (2021).

88. Costa, L. B. et al. Insights on SARS-CoV-2 Molecular Interactions With the Renin-Angiotensin System. Front. Cell Dev. Biol. 8, 559841 (2020).

89. Park, B. K. et al. Differential Signaling and Virus Production in Calu-3 Cells and Vero Cells upon SARS-CoV-2 Infection. Biomol Ther (Seoul) 29, 273–281 (2021).

90. Liao, M. et al. Single-cell landscape of bronchoalveolar immune cells in patients with COVID-19. Nat Med 26, 842–844 (2020).

91. Boumaza, A. et al. Monocytes and macrophages, targets of SARS-CoV-2: the clue for Covid-19 immunoparalysis. http://biorxiv.org/lookup/doi/10.1101/2020.09.17.300996 (2020) doi:10.1101/2020.09.17.300996.

92. Ahn, J. H. et al. Nasal ciliated cells are primary targets for SARS-CoV-2 replication in the early stage of COVID-19. Journal of Clinical Investigation 131, e148517 (2021).

93. Wang, S.-C. et al. Tannic acid suppresses SARS-CoV-2 as a dual inhibitor of the viral main protease and the cellular TMPRSS2 protease. Am J Cancer Res 10, 4538–4546 (2020).

